# Kinetics and Mapping of Ca-driven Calmodulin conformations on Skeletal and Cardiac Muscle Ryanodine Receptors

**DOI:** 10.1101/2023.07.13.548909

**Authors:** Robyn T. Rebbeck, Bengt Svensson, Jingyan Zhang, David D. Thomas, Donald M. Bers, Razvan L. Cornea

**Author notes:** Correspondence: Razvan L. Cornea, Department of Biochemistry, Molecular Biology, and Biophysics, University of Minnesota, 321 Church Street SE, Minneapolis, MN 55455, USA. Both authors contributed equally to this work (Robyn T. Rebbeck, Bengt Svensson).

## Abstract

Calmodulin (CaM) transduces [Ca^2+^] information regulating the rhythmic Ca^2+^ cycling between the sarcoplasmic reticulum and cytoplasm during relaxation and contraction in cardiac and skeletal muscle. However, the structural dynamics by which CaM modulates the SR Ca^2+^ release channel (ryanodine receptor, RyR) at physiologically relevant [Ca^2+^] is unknown. Using fluorescence lifetime detection of FRET between RyR-bound FKBP and CaM, we resolved different structural states of CaM and Ca-driven shifts in the conformation of CaM bound to RyR. We found that CaM becomes more compact in contracting high-Ca^2+^ vs. relaxing low-Ca^2+^. Skeletal and cardiac RyR isoforms show different CaM-RyR conformations, and binding and structural kinetics reflect functional roles. Furthermore, our FRET methods provide insight representative of physiological CaM-RyR structure, in synergy and comparison with cryo-EM models, which result from more disrupted samples. This technology will drive future studies focusing on pathologic CaM-RyR interactions, and on RyR dynamics with other important modulators.

## INTRODUCTION

Calcium cycling between the sarcoplasmic reticulum (SR) and the sarcoplasm is essential for muscle contraction and relaxation, which are primarily maintained by ryanodine receptors (RyR), and SR/ER Ca^2+^-ATPases (SERCA), respectively. Of the three mammalian RyR isoforms, RyR1 and RyR2 dominantly represent RyR in skeletal and cardiac muscle, respectively. The ∼65% sequence identity between isoforms is consistent with the overall structural similarity of the large (2.3 MDa) homotetrameric state observed via cryo-EM. Within the last decade, cryo-EM studies have greatly enhanced our understanding of RyR structural states, resulting in near-atomic resolution structures (3.2-4.2Å) of both RyR1 and RyR2 ^1, 2, 3, 4, 5, 6, 7, 8, 9, 10, 11, 12^. Although cryo-EM represents a powerful approach that has led to major breakthroughs, it comes with the inherent caveat that it resolves the highly populated static structures, and the full RyR sequence has yet to be charted. In addition to RyR structures, cryo-EM has been used to reveal RyR co-structures with its important regulators, e.g., calmodulin (CaM)^3, 4^. CaM-mediated regulation is particularly important for RyR2 proper function, where dysregulation has been associated with cardiac hypertrophy, arrhythmia and heart failure ^13, 14, 15, 16, 17, 18, 19^. Indeed, enhancing the CaM-RyR2 interaction has been shown to restore healthy Ca^2+^ cycling in cardiomyocytes ^20, 21^. Therefore, it is of key translational relevance to understand the mechanisms of long-range allosteric regulation of the channel opening by modulators, including CaM, that bind to the large cytosolic portion of the RyRs.

Calmodulin is a 17-kDa protein that contains four EF-hands between two terminal domains (known as the N- and C-lobe). These Ca^2+^ binding domains enable finely tuned modulation of RyR during Ca^2+^ cycling. Both Apo- and Ca^2+^-bound forms of CaM have been shown to bind to RyR1 and RyR2 with a stoichiometry of four CaM per channel ^22, 23^. Ca^2+^ binding affinity (KCa) to CaM has been shown to be enhanced from ∼3 to ∼1 µM by binding to RyR1 in SR membranes ^24^, providing Ca^2+^ sensitivity more in tune with Ca^2+^ cycling during striated muscle contraction. CaM is a well-established inhibitor of both RyR1 and RyR2 at contracting (µM) [Ca^2+^] ^25, 26, 27, 28^, thought to contribute to termination of SR Ca^2+^ release preceding cellular relaxation^17, 29^. CaM also inhibits RyR2 at resting (nM) [Ca^2+^] ^27, 28^, which is considered protection from chronic SR Ca^2+^ leak during diastole^30^. In contrast, CaM increases RyR1 activity at resting [Ca^2+^] ^25, 26^, which is thought to prime the channels for activation during EC coupling^29^. Based on low-resolution cryo-EM maps^31, 32, 33^, this divergence in functional effect has been previously suggested to reflect a difference in the CaM binding site on RyR1 vs. RyR2 in resting [Ca^2+^]. Specifically, cryo-EM mapping of apo-CaM to RyR2 assigned its binding to a site that is more similar to Ca^2+^-CaM (than apo-CaM) binding site on RyR1^33^. However, higher resolution cryo-EM structures (Fig. 1) place apo-CaM on RyR2 in a similar position as the high-resolution structures of apo-CaM binding to RyR1^3^, with the N-lobe interfacing with three surfaces in the RyR2 Helical Domain 1, and C-lobe interfacing with two surfaces in the cleft between the Central and Handle domains^4, 5^. The RyR sequences at the apo-CaM binding interfaces appear largely conserved between RyR1 and RyR2 ^3, 4^, which suggests that the factor mediating the isoform-specific functional effect of CaM is a sequence difference between RyR1 and RyR2 that allosterically controls the functional effect of CaM in low Ca^2+^. Without a CaM-RyR structure at physiologically low Ca levels, it may be difficult to elucidate the cause of isoform specific regulation by apo-CaM, as apo-CaM may not represent the structural state of CaM bound to RyR2 in nanomolar Ca^2+^. In the Ca^2+^-bound form, CaM’s N-lobe shifts down on RyR2 to bind wrapping around the canonical CaM binding site on RyR2 between the Helical Domain 2 and the Central domain^4^, which was supported by a prior crystal structure of Ca-CaM binding to RyR peptide (corresponding to RyR1 aa 3614–3643) ^34^.

**Fig. 1.**
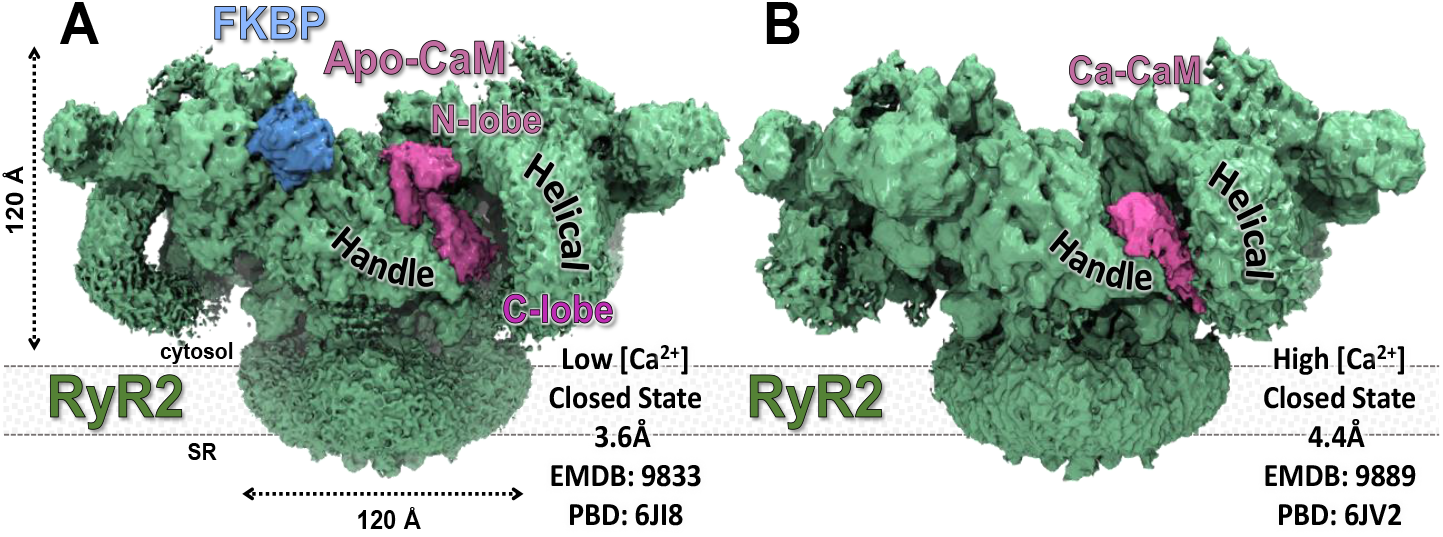
Cryo-EM maps of RyR2 reveal CaM binding to RyR2. Ca-free (apo) and Ca-bound -states of CaM bind at distinct locations. A) RyR2 in the closed state with FKBP12.6 (Light Blue) and apo-CaM (Purple) in low [Ca^2+^]^**4**^. B) RyR2 in the closed state with Ca-CaM bound at 20μM [Ca^2+^].

We have previously used steady-state FRET to resolve the orientation of CaM lobes relative to a fluorescent probe bound to FKBP12.6 on RyR^35, 36^. These measurements have been used to map the position of divergent regions of the RyR1^37^, and the RyR2 binding site of DPc10, a peptide corresponding to RyR2 2460-2495^38^. A limitation to this approach is that steady-state fluorescence measurements of FRET yield a single distance for the ensemble of observed donor-acceptor distances. To overcome this shortcoming, we have used fluorescence lifetime (FLT) measurements of FRET, which can be analyzed to resolve up to three populations of donors: two at measurable distances from the acceptor, and one at a distance outside the FRET range (>100 Å)^39, 40^. Using this technology to resolve distances between 8 distinct probe sites on FKBP12.6 and 4 distinct probe sites on CaM when bound to RyR1 and RyR2 in SR membranes, we have refined CaM binding and advanced our methodology for mapping the binding sites of many RyR modulators. Overall, these FRET-based structural models can be used to suggest whether the cryo-EM models likely represent the CaM-RyR state in a more physiological environment. Further, we adapted this methodology for the first measurements of CaM association to RyR1 vs RyR2 and kinetics of Ca^2+^-driven structural transition in CaM bound to RyR, within the time scale of muscle contraction.

## RESULTS

### Labeling FKBP12.6 and CaM does not alter binding to RyR

For labeling CaM and FKBP12.6, we substituted single cysteines at sequence locations in CaM (no native Cys) and Cys-Null FKBP12.6^41^ that would be solvent exposed and unlikely to interfere with RyR interaction (see next section in Results). We have previously evaluated the binding and functional effect on RyR1 of labeling FKBP12.6 with Alexa Fluor 488 C5 maleimide (AF488) at sites T14C, E32C, R49C and T85C^35^, and labeling CaM with Alexa Fluor 568 C5 maleimide (AF568) at sites T34C and T110C^41^. Here we extended our collection of single-Cys substitutions to FKBP12.6 sites G1, T6, K44 and Q65 (Supplementary Fig. 1A), and CaM sites T26 and Y99 (Supplementary Fig. 2A). Thus, we used eight AF488-labeled FKBP12.6 single-Cys mutants and four AF568-labeled CaM single-Cys mutants.

To confirm that mutation and attachment of a fluorescent probe does not alter the necessary high affinity of FKBP12.6 binding to RyR, we monitored co-sedimentation of each AF488-FKBP12.6 variant with RyR1 in skeletal SR membranes. The variants (T14C, E32C, R49C and T85C) that were previously shown to bind similarly with WT-FKBP12.6^35^, displayed very similar binding affinities (3.7-5.8 nM K_d_ values) for RyR1 (Supplementary Fig. 1C). Of the new mutants, K44C and Q65C displayed affinities within this range, and G1C and T6C showed slightly lower affinities (12.6 and 7.3 nM, respectively) (Supplementary Fig. 1C). Overall, these high affinities are suitable for our FRET measurements, and demonstrate that labeling did not disrupt binding of FKBP12.6 to RyR. With the binding of FKBP12.6 very similar on RyR1 and RyR2, it is reasonable to assume that, like RyR1, labeling at any of these eight sites does not impact FKBP12.6-RyR2 binding.

To determine the impact of fluorescence labeling of Ca-sensitive and Ca-insensitive CaM on RyR interaction, we used [^3^H]ryanodine binding measurements, as previously^41^. Using the level of [^3^H]ryanodine binding as an index of channel activity, we show that addition of 800 nM unlabeled WT-CaM significantly increases RyR1 activity at 30 nM Ca^2+^, and decreases RyR1 activity at 30 µM Ca^2+^ (Supplementary Fig. 2C). In alignment with Ca-insensitivity, addition of 800 nM unlabeled Ca-insensitive CaM (CaM_1234_) increased RyR1 activity at both 30 nM and 30 µM Ca^2+^ (Supplementary Fig. 2C). The functional effects of unlabeled CaM align with previous publications^25, 28, 40, 41^. As shown in Supplementary Fig. 2C and D, the Cys substitution and labeling of CaM does not impact the direction of functional effect at most sites. Although, the functional effect of CaM on RyR1 is moderately increased by labeling at site T110 (Supplementary Fig. 2C). In alignment with previous studies, we show that unlabeled WT-CaM decreases RyR2 activity at 30 nM and 30 µM Ca^2+^, which is significantly lost in CaM_1234_ (Supplementary Fig. 2D). Similar to the functional effect on RyR1, labeling at sites T26, T34 and Y99 did not impact the direction of functional effect (Supplementary Fig. 2). However, labeling at site T110 appears to deviate from unlabeled constructs, which is most obvious in comparison with WT-CaM at 30 nM Ca^2+^, where AF568-T110C-CaM increased RyR2 activity (Supplementary Fig. 2D). Our functional assays indicate that the F-CaM mostly retained functional interactions with RyR1 channels that are characteristic of WT CaM at nM and µM Ca^2+^.

### Effective positions of the FKBP- and CaM-attached probes

To interpret the FKBP-CaM FRET data, we used an *in silico* approach to determine the location of the donor probes attached to FKBP in RyR1 and RyR2. For the computational studies of RyR1, the structure chosen was the 3.6-Å closed state and the corresponding cryo-EM density map (Supplementary Fig. 7)^1^. For RyR2, we used the 3.6-Å structure corresponding to the closed state and the corresponding cryo-EM density map (Fig. 2, Supplementary Fig. 3)^4^. We chose the closed structures because they reflect the more physiological state that predominates in our FRET samples.

**Fig. 2.**
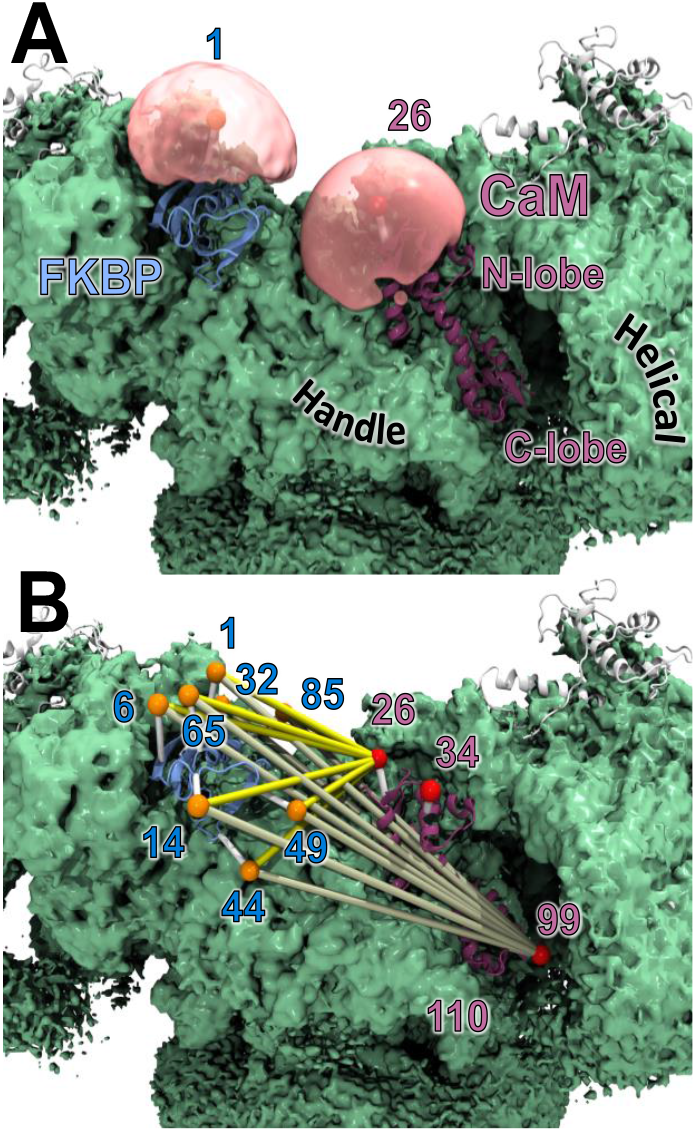
Labeling and locations of FRET probes. In silico predictions of the donor and acceptor fluorophore locations were conducted by simulated annealing for donors bound to FKBP, and by a method based on probability distributions for acceptors bound to CaM. For these calculations, we used cryo-EM structures of RyR1 and RyR2 in closed states, at both high and low Ca^2+^. Shown here are results for RyR2 at low Ca^2+^ (EMDB: 9833, PDB: 6JI8)^**4**^. A) The volume sampled by the fluorophore is shown as a pink bubble for the donor attached at FKBP residue 1. The average center-position of the probe is shown as an orange sphere inside the pink bubble. The exemplar CaM probe location is displayed as a pink bubble with the average acceptor location shown as a red sphere. B) The predicted FRET distances from all donor-labeled sites on FKBP to acceptor-labeled site 26 in the N-lobe and site 99 in the C-lobe of CaM, are shown with yellow and tan colored lines, respectively. The acceptor probe bound at CaM residue 110 is hidden behind the handle domain.

In the atomic structure of RyR1 and RyR2, the eight residues at positions 1, 6, 14, 32, 44, 49, 65, and 85 of FKBP12.6 that were used for probe attachment in the fluorescence experiments were mutated to cysteine *in silico*. The AF488 probe was attached to each of the eight cysteines, generating eight starting conformations for the simulated annealing calculations used to determine the effective donor probe coordinates. For comparison with our trilateration and to calculate predicted distances between donor and acceptor probes in the cryo-EM structure, we used a new faster method to determine the region in space the probe can sample and the average probe location on CaM at sites 26, 34, 99, and 110, as described in *Methods*.

### Resolving distance distribution between probes from FLT-FRET experiments

To map the position of CaM in 30 nM and µM Ca^2+^, we used FLT-FRET acquired using time correlated single photon counting (TCSPC), which can be analyzed to yield distance relationships between the established donor probe sites on FKBP12.6 and acceptor probe sites on CaM (e.g., as in Fig. 3B). In each experiment, we used SR membranes from porcine skeletal or cardiac muscle that were labeled with one of the eight D-FKBP variants validated above. FLT waveforms (e.g., as in Fig. 3A) were acquired 2 hrs after incubation of our FRET samples with A-CaM or A-CaM, both with 4 variations by being labeled at one of four sites, as described above. We have previously used single distances calculated from FRET efficiencies, to locate acceptor-labeled sites on RyR cryo-EM maps^37, 38^. Here, we enhance this method, as FLT-FRET allows for simultaneous detection of CaM and FKBP12.6 binding to RyR and structural analysis. As previously used, this allows us to integrate the fraction of unbound donors (∼10%; not participating in FRET) in the distance calculations^40, 41^. Furthermore, with the possibility of FRET from neighboring faces of the tetrameric RyR complex, extracting multiple-distance information enhances the resolution of the actual distance relationships between D-FKBP and A-CaM bound on the same face of RyR.

**Fig. 3.**
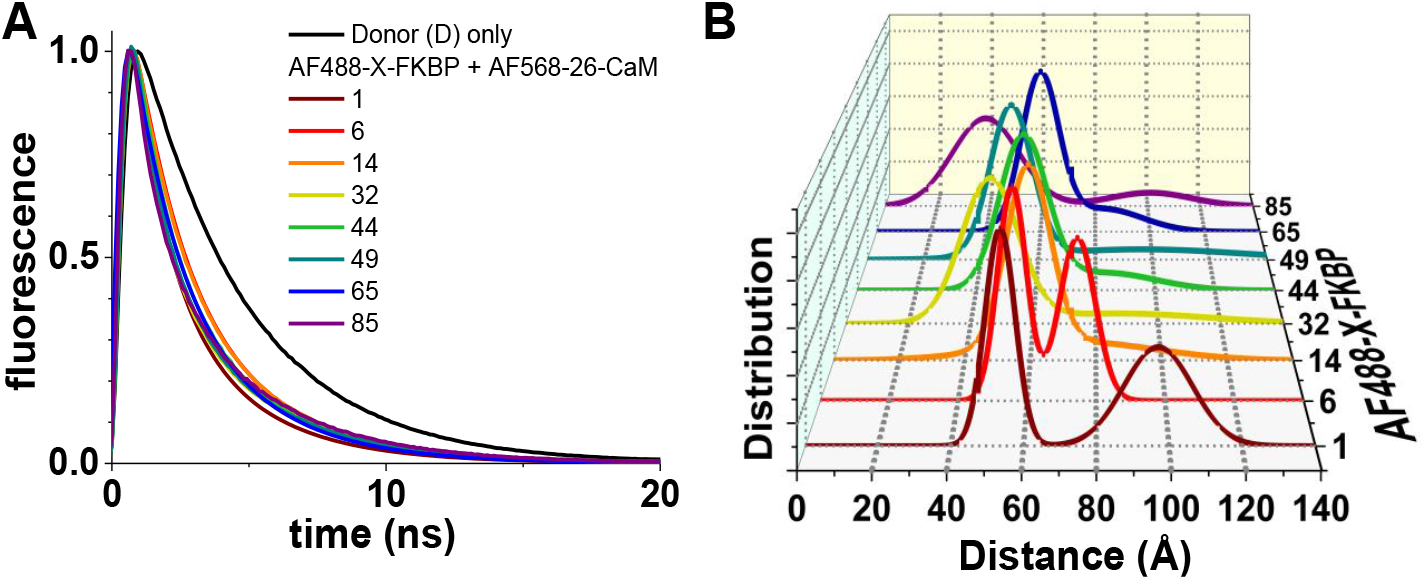
Representative FLT-FRET detection of distance distributions between D-FKBP and A-CaM probes. At 30 nM Ca^2+^, SR membranes from skeletal muscle were labeled with D-FKBP (AF488-X-FKBP; X = Cys mutation site for fluorescence labeling), and then incubated with 800 nM CaM labeled with acceptor probe at the N-lobe residue T26C. A) Representative FLT-detected waveforms for D-labeled at 8 different sites. B) Multiexponential analysis of FLT-FRET data yielded a two–distance Gaussian distribution model for the separation between D-FKBP and A-CaM within RyR1. Averaged data shown in Supplementary Fig. 3.

With the CaM N-lobe probe locations being closer to probes on FKBP12.6 (Fig. 1), a two-distance Gaussian distribution model of separation between D-FKBP and A-CaM represented the best fit of the FLT-FRET data (Fig. 3). However, the longer distances between D-FKBP and CaM C-lobe probe locations produced only a single-distance Gaussian distribution. This aligns with our previous report using the same model with AF488-R49C-FKBP12.6 and N-lobe labeled CaM (T34C) vs C-lobe labeled CaM (T110C)^40^. The average features of each Gaussian fit are shown in Supplementary Fig. 3-6. This type of data was used for trilateration, as shown in Supplementary Table 1-4.

### Trilateration of CaM binding on RyR 3D map demonstrates Ca^2+^-driven structural shifts

The method of using distances to determine a location in space is termed trilateration. This method is most commonly known for its use in the Global Positioning System (GPS), where known satellite-positions are used to determine an antenna location. The use of trilateration based on FRET distances and macromolecules has been previously described by us and others^38, 42, 43^. From the simulated-annealing calculations above, we obtained coordinates of the effective donor locations, which we used for trilateration. We used effective AF488 locations calculated for each of the eight FKBP attachment sites (1, 6, 14, 32, 44, 49, 65, and 85) together with Gaussian distance distributions calculated from FLT-FRET analysis, to determine a locus in space corresponding to acceptors attached at each of the four locations on CaM (26, 34, 99, and 110). In most cases, the shortest distance was the most populated, but there were a few cases where the longer distance was more populated. Upon comparison of acceptor loci sites, we chose to trilaterate using the shortest distance, except when the longer-distance population was more than double the shorter distance population (Supplementary Fig. 8).

As expected, all acceptors were localized to the grove between the handle and helical domain on RyR, with the CaM N-lobe oriented above its C-lobe (Fig. 1 and 4). To differentiate between Ca^2+^-driven structural shifts in RyR vs CaM, we compared the loci of probes bound to CaM_1234_ in nM and µM Ca^2+^. The acceptor loci for CaM_1234_ did not shift between nM and µM Ca^2+^, suggesting that any Ca^2+^-driven shift observed with probes bound to normal CaM is due to its own structural shift, and not of RyR.

**Fig. 4.**
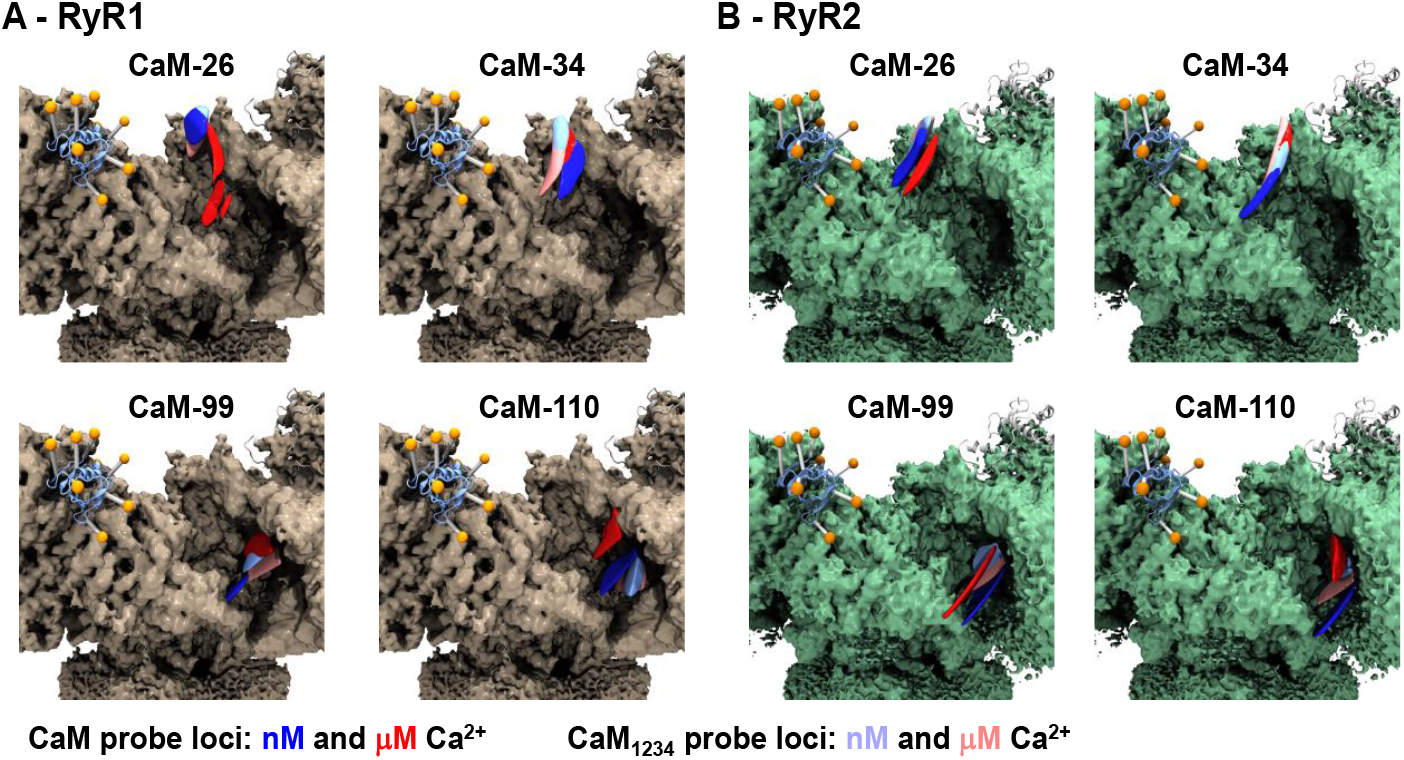
Location of trilaterated acceptor fluorophore loci on RyR1 and RyR2. Cryo-EM maps of A) RyR1 (EMDB: 8342, PDB: 5T15)^1^ and B) RyR2 (EMDB: 9833, PDB: 6JI8)^4^ bound to the atomic structure of FKBP (orange), with trilaterated loci of AF568 attached to CaM residues as indicated. Trilaterated loci of probes bound to CaM in assay conditions containing 30 nM and 30 µM free Ca^2+^ are represented in blue and red, respectively. Trilaterated loci for probes bound to Ca^2+^ insensitive CaM (CaM_1234_) in assay conditions containing 30 nM and 30 µM free Ca^2+^ are represented in light blue and light red, respectively.

The probe locus of AF568-26C-CaM on RyR1 was driven down and outwards by the nM to µM shift in Ca^2+^ (Fig. 4A and Supplementary Fig. 9). The probe locus of AF568-34C-CaM (also in the N-lobe) did not detectably shift between nM and µM [Ca^2+^] for both CaM and CaM_1234_ (Fig. 4A and Supplementary Fig. 9). Together, these suggest that Ca^2+^ binding to CaM structurally results in a rotation of the N-lobe as it shifts downwards and out within the RyR1 binding groove. Similar to the probe at residue 34, the probe at CaM residue 99 showed a small shift for both CaM and CaM_1234_ (Fig. 4A and Supplementary Fig. 9). In contrast, the probe locus of AF568-110C-CaM was distinctly shifted upward and inward in µM Ca^2+^ relative to nM Ca^2+^. For both residue 26 and 110 acceptor attachment sites in CaM, the loci in nM Ca^2+^ overlap with the corresponding loci of CaM_1234_ at both nM and µM Ca^2+^ (Fig. 4A and Supplementary Fig. 9). This suggests that at nM Ca^2+^, both CaM lobes reflect an apo (Ca^2+^ unbound) structural state. Overall, the nM to µM Ca^2+^ shift appears to drive RyR1-bound CaM to form a more compact state, with the lobes rotating in opposite directions and driving closer together.

The probe locus of AF568-26C-CaM on RyR2 displays a similar Ca^2+^ driven downward and outward shift as we observed with RyR1, although more subtle. However, at nM Ca^2+^ this locus in normal CaM is distinctly positioned between the equivalent locus in CaM_1234_ and the µM Ca^2+^ locus (Fig. 4B and Supplementary Fig. 9). This positioning suggests at least partial Ca^2+^ occupancy of the CaM N-lobe at nM Ca^2+^. In contrast to RyR1-bound CaM, the probe locus of AF568-34C-CaM in RyR2 is similar between normal CaM and CaM_1234_, but distinctly shifted lower in nM relative to µM Ca^2+^ (Fig. 4B and Supplementary Fig. 9). Acceptor loci at both CaM-99 and CaM-110 display an upward shift due to nM to µM [Ca^2+^]. Additionally, the CaM-110 probe locus appears to shift inward between nM and µM Ca^2+^ (Fig. 4B and Supplementary Fig. 9). Similar to RyR1-bound CaM, the nM to µM Ca^2+^ shift appears to drive RyR2-bound CaM to form a more compact state, with the lobes rotating in opposite directions and moving closer together. In contrast to RyR1-bound CaM, CaM bound to RyR2 in nM Ca^2+^ may be in a structural state with partial occupation by Ca^2+^ binding.

### Kinetics of CaM-RyR binding using stopped-flow FLT-FRET

An advantage of direct waveform recorded FLT-FRET technology is that it is compatible with stopped-flow kinetic studies, as we previously reported with myosin-focused, transient structure studies^44, 45^. Armed with the probe pair that displayed the largest distance difference between nM and µM Ca, D-85-FKBP and A-26-CaM (Supplementary Table 1 and 3), we carried out stopped-flow FLT-FRET measurements to first investigate CaM binding. Our intent was to explore the capabilities of this assay and technology. With 2-ms mixing time and 0.2 ms acquisition rate, this technology enabled the first measurements of CaM-RyR association kinetics, and a direct comparison between the skeletal and cardiac RyR isoforms from porcine tissue. This complements the field’s current knowledge of full-length RyR isoform (RyR1 vs RyR2) impact on CaM dissociation rates, specifically with Meissner and colleagues (2009) demonstrating that CaM dissociates from canine RyR2 three-fold faster relative to rabbit RyR1 in 150 nM Ca^2+ 46^.

As above with our trilateration studies, to resolve differences between apo- and Ca^2+^-CaM, we concurrently monitored binding of CaM_1234_ to RyR1 and RyR2 in 30 nM and µM Ca^2+^. In all conditions, the time courses are best fit using three exponentials (Fig. 5 and Supplementary Fig. 12 and 13). As shown in Fig. 5, the amplitude-weighted time constants for RyR2 are 3.3- and 6-fold faster than RyR1 at 30 nM and µM Ca^2+^, respectively. These results indicate that CaM-RyR1 association kinetics does not significantly differ between relaxing (nM Ca^2+^) and contracting (µM Ca^2+^) conditions (Fig. 5). The CaM_1234_-RyR1 association kinetics under both Ca^2+^ conditions was similar to that of CaM-RyR1 (Supplementary Fig. 13). In contrast, the CaM-RyR2 association was significantly (∼2-fold) faster at 30 µM Ca^2+^ relative to CaM at 30 nM Ca^2+^ and CaM_1234_ at both Ca^2+^ (Supplementary Fig. 13). Overall, this reflects that CaM binds faster to RyR2 vs RyR1, and that binding is [Ca^2+^] sensitive for RyR2 but not for RyR1.

**Fig. 5.**
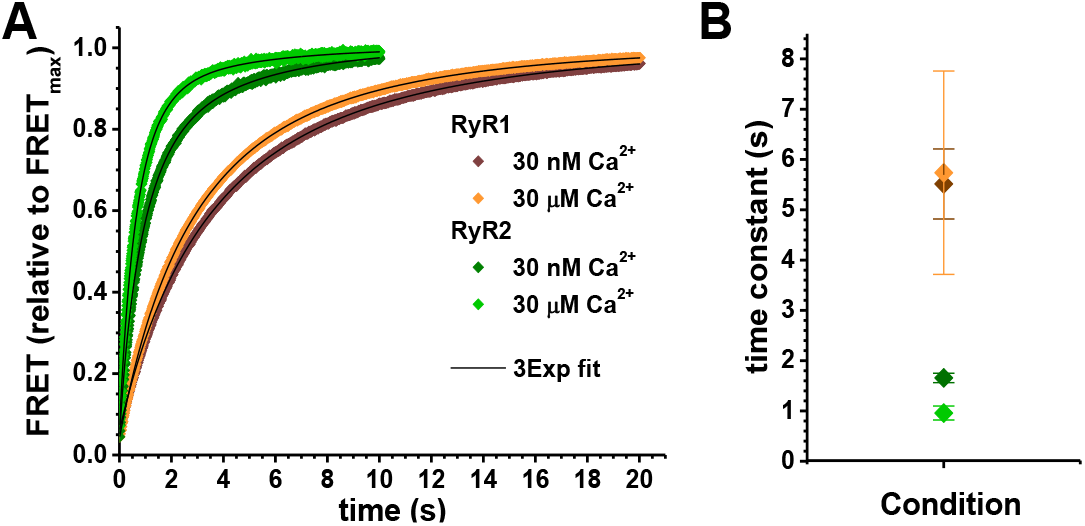
Kinetics of A-CaM binding to RyR1 and RyR2 in 30 nM and µM Ca^2+^. SR membranes from porcine skeletal (brown) or cardiac (green) muscle were labeled with D-FKBP (AF488-85-FKBP), and then FLT time course was acquired after rapid (2 ms) mixing with 800 nM (final) A-CaM (AF568-26-CaM. A) Representative FLT time course following mixing. B) Amplitude-weighted average time constant from three-exponential analysis of FLT-FRET data for CaM binding to RyR1 or RyR2 at 30nM or 30 µM Ca^2+^. Data shown as mean±SEM, n = 3. Parameters from three-component fitting are shown in Supplementary Fig. 13.

### Kinetics of Ca^2+^-driven structural shift of RyR bound CaM

With distinct Ca^2+^-dependent distances between donor bound to 85-FKBP and acceptor 26-CaM (Supplementary Table 1 and 3), we proceeded to use this probe pair to observe the kinetics of [Ca^2+^]-driven A-CaM structural transitions. Specifically, focusing on the 30 nM to µM and 30 µM to nM Ca^2+^ transitions. For A-CaM bound to RyR1, we show that the structural transition is faster (3.4-fold) going from low (nM) to high (µM) Ca^2+^, relative to the high to low Ca^2+^ transition (Fig. 6A,B). For A-CaM bound to RyR2, the high to low Ca^2+^-driven structural transition, is 4-fold slower (p = 0.025) relative to A-CaM bound to RyR1 (Fig. 6).

**Fig. 6.**
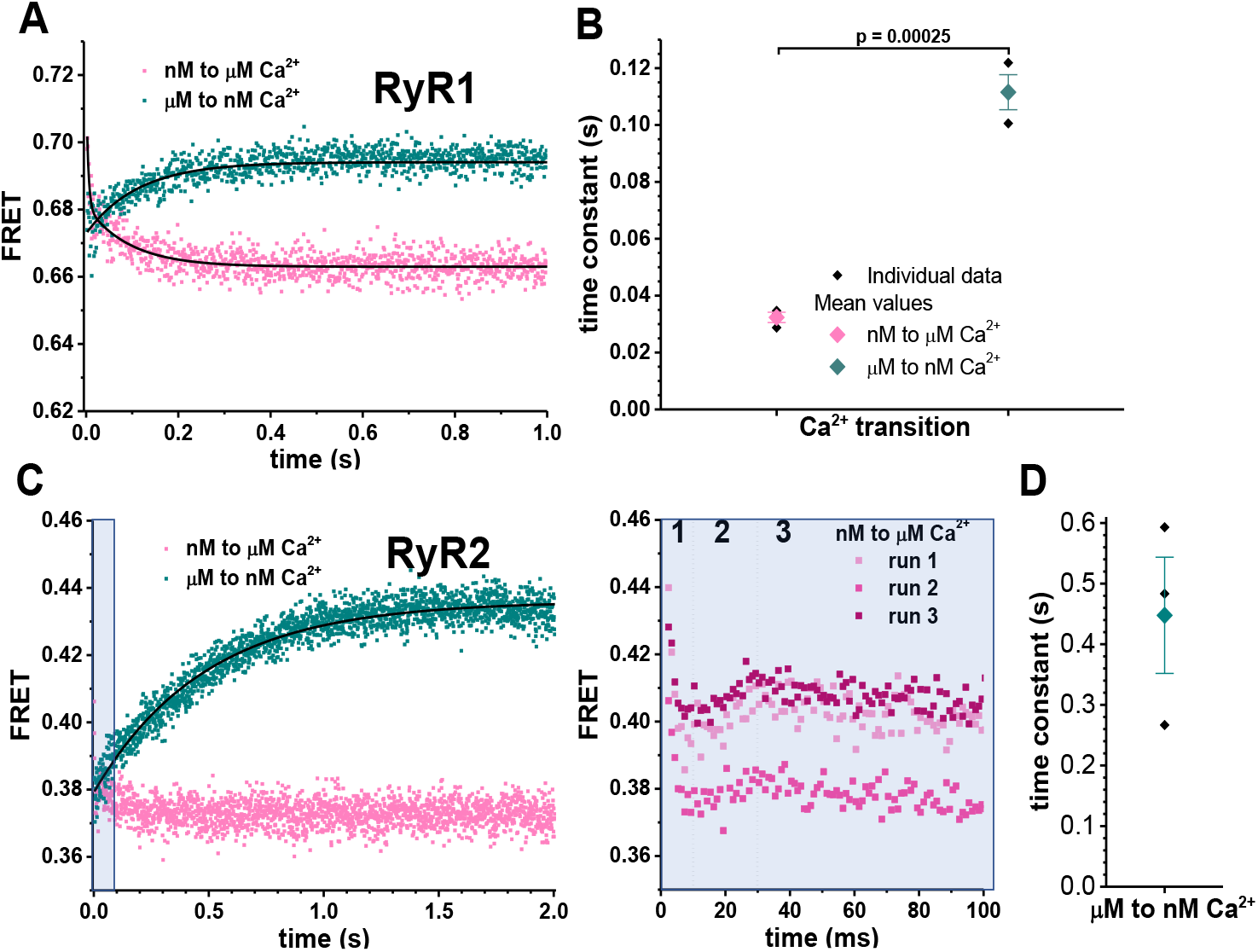
Calcium-driven structural transitions of A-CaM bound to RyR1 or RyR2. SR membranes from porcine skeletal and cardiac muscle were labeled with D-FKBP (AF488-85-FKBP), incubated with 1.6 µM CaM and then FLT time course was acquired after rapid (2 ms) mixing with Ca^2+^ to increase [Ca^2+^] from 30 nM to µM (pink), or EGTA to decrease [Ca^2+^] from 30 µM to nM (green). Representative FLT waveforms shown in Supplementary Fig. 14. A) With A-CaM bound to RyR1, representative FLT-FRET time course following rapid [Ca^2+^] shift. FRET data for nM to µM Ca^2+^ transition fit best to two-exponential analysis with average parameters shown in Supplementary Fig. 15. For µM to nM, Ca^2+^ transition fit best to one-exponential analysis. B) Averaged time constants per Ca^2+^ transition. Data shown as mean ± SEM, n = 3. Significance calculated using unpaired, two-way Student’s T-test. C) With A-CaM bound to RyR2, representative FLT-FRET time courses following rapid [Ca^2+^] shift. For µM to nM, Ca^2+^ transition fit best to one-exponential fitting. Within the first tenth of a second, FRET data for nM to µM Ca^2+^ transition was multiphasic, which was shown to be reproducible, as illustrated in the right panel, where each phase 10ms, 20-30ms and > 30 ms is indicated by numbers 1, 2 and 3, respectively. D) Time constant for µM to nM Ca^2+^ transition for A-CaM bound to RyR2.

With the 30 nM to µM Ca^2+^ transition, A-CaM bound to RyR2 undergoes a multiphase shift in FRET (Fig. 6C, right panel). FRET drops rapidly within the first 10 ms (stage 1), followed by a FRET increase from 10 to 20 ms, and ending with a slower FRET decrease beyond 30 ms (Fig. 6C, right panel). With saturating A-CaM bound, and slower association kinetics, these Ca^2+^-dependent FRET changes might reflect stepwise structural changes in the RyR2-bound CaM.

RyR isoform comparison for the nM to µM Ca^2+^ driven shift in bound CaM shows that stage 1 and stage 3 of CaM-RyR2 kinetics is somewhat reflected in the CaM-RyR1 kinetics, with 4.2 and 78.6 ms time constants from the best fit two-exponential analysis (Supplementary Fig. 15). Stage 2 is particular to RyR2.

## DISCUSSION

We have advanced our FRET-based technology for molecular trilateration to map Ca^2+^-driven structural transitions of RyR-bound CaM in native SR membranes, under physiologically relevant conditions. Furthermore, we have exploited rapid FLT data acquisition – as enabled by direct-wave recording technology implemented in a stopped-flow apparatus – for breakthrough observations of this structural transition’s kinetics at a millisecond time-resolution. Existing cryo-EM maps provide important constraints for our analyses. Although extremely informative for the overall understanding of the RyRs, the cryo-EM reconstructions come with the significant caveat that they describe static snapshots of the most populated structural states of the RyR channel complex, after removal from its physiological environment. Our initial stopped-flow kinetics observations provide unique insight into the association of CaM to RyR1 vs RyR2 – at a physiologically relevant time-scale – thus completing the description of CaM-RyR binding kinetics that had started with previous studies of the much slower CaM dissociation from RyR1 vs RyR2^46^.

### CaM binding in the RyR1 and RyR2 cryo-EM structure

The first high-resolution cryo-EM structures published with CaM bound to RyR were for RyR2^4^. CaM was observed to be bound between the Handle and the Helical domains in all six structures published for RyR2. These structures are from different conditions that are either in the closed or open state of RyR2. Here we are interested in the state of RyR that predominates under the physiological conditions of our experiments, i.e., the closed state in both low and high Ca^2+^. The open state is present only briefly when ATP is not present. Apo-CaM on the closed state, as represented by structure 6JI8, is bound high in the groove between the Handle and Helical domains. The N-lobe is positioned high in this groove and makes contact with both of these RyR2 domains. The C-lobe is positioned lower in the groove making contact mostly with the Handle domain, but also with the C-terminal portion of the α-helix of RyR2 (aa 3593-3606) which is part of CaM Binding Domain 2 (CaMBD2). Helical Domain 2 precedes the CaMBD2 region in the RyR2 sequence (and would fill the lower portion of this groove). However, Helical Domain 2 is mostly unresolved in these RyR2 structures. Ca-CaM in the closed state of RyR2, as represented by structure 6JV2, is bound further down in the groove, and with the two lobes of CaM now wrapped around the CaMBD2 α-helix (aa 3585-3606). The N-lobe is rotated about 180° and shifted lower in the groove compared with the apo-CaM structure. The C-lobe has undergone a rotation and a shift towards Helical Domain 2.

For RyR1 there are two available cryo-EM structures with CaM bound. One is with apo-CaM^3^ and the other with Ca-CaM^47^. Both these structures are in the closed state of the RyR1 channel. The apo-CaM structure has FKBP12.6 bound while the Ca-CaM has FKBP12 bound. Apo-CaM, as represented by structure 6×32, is bound in the same location as previously described for RyR2, i.e., in the groove between the Handle and Helical domains with the N-lobe positioned high in this groove and making contact between these two RyR domains. The C-lobe is positioned lower in the groove, making contact mostly with the Handle Domain, but also with C-terminal portions of the CaMBD2 region of RyR1 (aa 3625-3634). Helical Domain 2 is better resolved in RyR1, but it does not make any contact with apo-CaM. Ca-CaM in RyR1, as represented by structure 7TZC, is bound differently compared with Ca-CaM in the RyR2 structure. The N-lobe is bound high in the groove in a similar location as apo-CaM. CaM helix I is in the same location as in the apo-CaM structure but helices II and III are rotated upward toward Helical Domain 1 by about 45°. CaM helix IV (which connects the two lobes) has rotated toward the RyR1 Helical Domains to allow the C-lobe a new binding location closer to Helical Domain 2. This is mostly a shift in binding location closer to Helical Domain 2 with just a minor rotation. In contrast to RyR2, in RyR1 only the C-lobe of Ca-CaM is in contact with CaMBD2 (aa 3614-3637).

The sample conditions are different in the RyR1 vs. RyR2 Ca-CaM bound structures. For the RyR2 Ca-CaM closed state structure, media include 20 µM Ca^2+^ and PCB95, and there is only partial binding of FKBP. Under these conditions the addition of Ca-CaM closes the channel that was activated by Ca^2+^ and PCB95^4^. For the RyR1 Ca-CaM closed state, the conditions were 30 μM Ca^2+^, 10 mM ATP, and 5mM caffeine, with FKBP12 present. Here the addition of Ca-CaM and also the Rycal drug ARM210 stabilizes the closed state as these conditions otherwise would generate the primed and open states. As the conditions are very different between the RyR1 and RyR2 Ca-CaM structures, it is not possible to conclude that the difference in binding of Ca-CaM is due to RyR isoform differences.

### FRET-trilaterated position of CaM within the RyR1 and RyR2 3D structure

By combining the trilaterated locations of the A-CaM probes (Fig. 4) we can determine the binding locations and get information on the orientation of the individual lobes of CaM at low and high Ca^2+^ in both RyR1 and RyR2. This allows us to directly compare the binding position of CaM in our experiments with the binding in the cryo-EM structures. At low Ca^2+^, FRET trilateration places the N-lobe of apo-CaM high in the groove between the Handle and Helical domains, with position 26 higher up than 34 in both RyR1 and RyR2 (Fig. 7). The C-lobe sites are bound lower in this groove, close to the Handle Domain. These are the same locations as shown in the cryo-EM structures for both RyR1 and RyR2 (compare with apo-CaM position of RyR2 in Fig. 1).

**Fig. 7.**
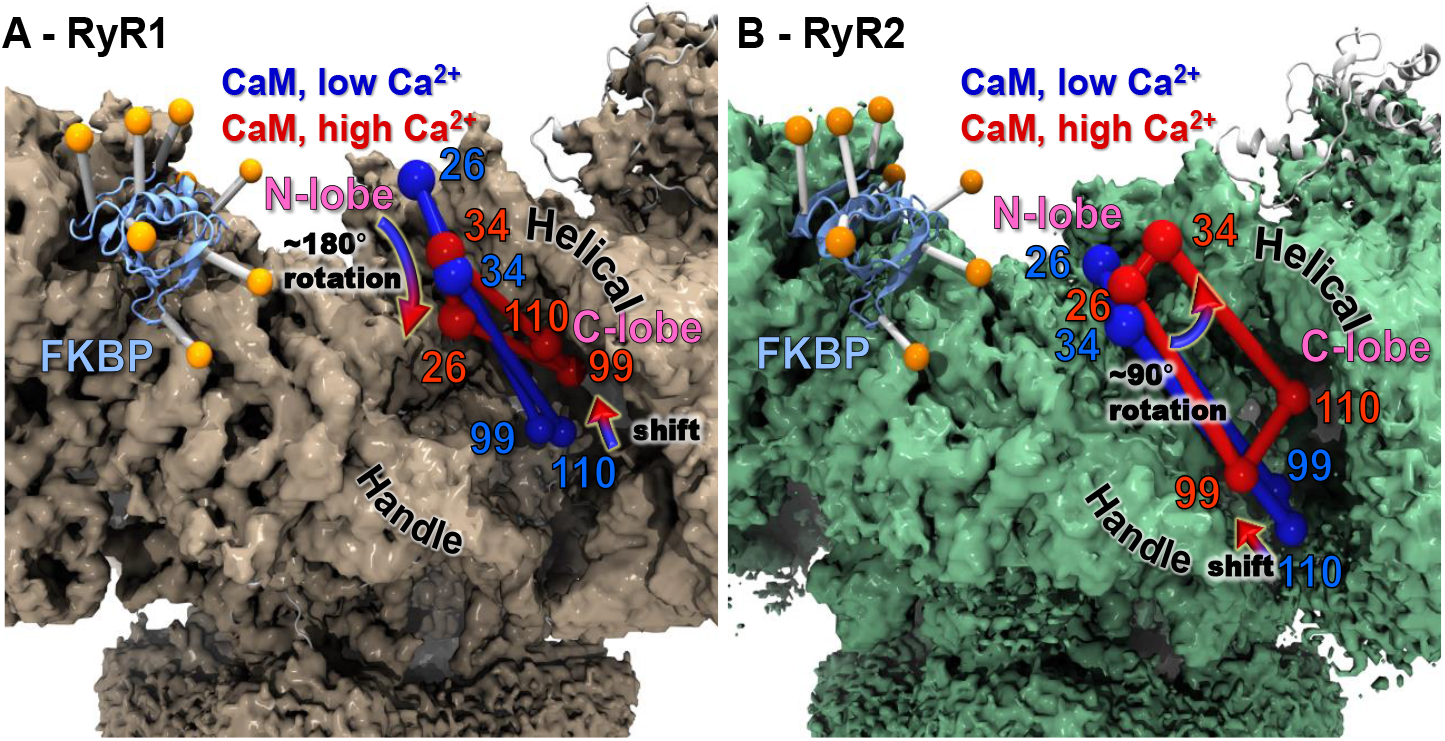
Overall Ca^2+^ driven structural transitions of A-CaM probe loci on RyR1 or RyR2. The center of the trilaterated loci for AF568 probes bound to indicated CaM residues are shown as spheres and drawn in blue for low Ca^2+^ (apo-CaM) and red for high Ca^2+^ (Ca-CaM). The spheres are connected by lines to better show the shape of and change in rotation of sites in the lobes of CaM. A) The cryo-EM density map of RyR1^**1**^ is shown in brown. FKBP is shown as ribbon representation in blue with the donor probe positions shown as orange spheres. A Ca^2+^ induced 180° rotation and a shift downward is observed in the N-lobe and a shift upward is observed in the C-lobe. B) The cryo-EM density map of RyR2^**4**^ is shown in green. FKBP is shown as ribbon representation in light blue with the donor probe positions shown as orange spheres. A Ca^2+^ induced ∼90° rotation is observed in the N-lobe and a minor shift upward is observed in the C-lobe. Arrows indicate direction and rotation of structural shift between low and high Ca^2+^.

In RyR1 at high Ca^2+^, we observe a large shift downward of position 26 while position 34 does shift minimally. This suggests a rotation of the N-lobe by about 180°. We also see a significant shift upwards of the C-lobe sites towards the Helical domains. Combined, these suggest that Ca-CaM is in a more compact conformation than apo-CaM. However, this is quite different from what is seen in the RyR1 Ca-CaM cryo-EM structure (7TZC). Our results with RyR1 are similar to what has been reported for Ca-CaM in the RyR2 cryo-EM structure (6JV2). This discrepancy between trilateration and cryo-EM suggests that there are more Ca-CaM RyR1 structural states that have not yet been detected by cryo-EM.

In RyR2 at high Ca^2+^, we observe a minor shift downward of position 26 and a shift towards the Helical domains for position 34. This suggests a rotation of the N-lobe by about 90°. We see an upward shift of the C-lobe sites, which is much smaller than what was observed in RyR1. Position 110 shifts closer to the Helical domains. Ca-CaM is becoming more compact than apo-CaM but not as compact as seen in RyR1. Therefore, the FRET trilateration is distinctly different from what is seen in the RyR2 Ca-CaM cryo-EM structure (6JV2), which has a very compact Ca-CaM bound lower in the groove. The trilaterated Ca-CaM binding location is closer to what is seen for Ca-CaM in the RyR1 cryo-EM structure (7TZC). However, our trilated results show CaM become more compact in response to Ca^2+^, which is not observed in the RyR1 cryo-EM structure. Our results suggest a conformation that is somewhere between the RyR1 and RyR2 cryo-EM structures. A possible interpretation of the trilateration results is that, under the more physiological conditions allowed by the FRET measurement, multiple locations of Ca-CaM coexist in the RyR2 CaM-binding site.

In conclusion our FRET trilateration can accurately locate the binding location and orientation of CaM in RyR. Our results match well with the position and orientation of CaM in the cryo-EM structures, especially for apo-CaM. Furthermore, our results suggest that there are different binding locations for Ca-CaM than reported in the current cryo-EM structures. One could speculate that if the sample conditions were the unified between cryo-EM and FRET, similar Ca-CaM binding would be detected. At this point, we do not have enough cryo-EM data to unambiguously describe the RyR isoform-specific differences in Ca-CaM binding. Our stopped-flow FRET kinetics results further support the existence of these differences.

### Characterizing CaM-RyR binding kinetics

Prior to this study, knowledge of the temporal relationships between CaM and RyR were limited, and mostly involved peptides corresponding to regions of RyR1/2. Here, we used a stopped flow apparatus with our FRET assay, to monitor the fast association rates of CaM binding to RyR1/2, as well as the faster rates of Ca^2+^-driven structural transition in CaM bound to RyR1/2. We provide evidence suggesting that the observed FRET changes are due to direct shifts in CaM binding to RyR through the CaM N-lobe, where the acceptor probe is bound. Due to low FRET, we were unable to resolve the kinetics of A-CaM binding and Ca-driven structural transition through monitoring FRET with the acceptor probe bound at the CaM C-lobe. This reduced resolution is probably due to the method of FLT data acquisition, as direct waveform recording was used for acquiring fluorescence decays in the kinetics experiments, whereas the more resolved but much slower FLT acquisition technology (time correlated single photon counting) was used for the acquisition of fluorescence decays used in trilateration (at equilibrium). An additional limitation of the kinetics measurements is that we could not confidently fit the Gaussian distance distributions to the data, as we did with the data used for trilateration.

Our results show that the CaM-RyR1 association kinetics does not shift between nM and µM Ca^2+^. This was surprising given that the functional effects of CaM on RyR1 differ so sharply between the two [Ca^2+^] tested.

In contrast, we show that CaM-RyR2 association kinetics is two-fold faster for µM relative to nM Ca^2+^, despite the inhibitory effect of CaM on RyR2 at both [Ca^2+^] tested. Notably, the level of CaM-mediated inhibition of RyR2 is greater at µM relative to nM Ca^2+^. In comparison between RyR1/2 isoforms, we also observed that a substantially faster CaM association with RyR2 relative to association with RyR1. This aligns with a previous report that CaM dissociation from RyR2 is three-fold faster than that of dissociation from RyR1^46^. In contrast to our observations with full-length RyR in native SR membrane preparations, a recent study using peptides corresponding to the highly conserved CaMBD2 of RyR1 (aa 3614-3640) and RyR2 (aa 3580-3608) showed that the rates of CaM association with RyR1 and RyR2 peptides did not significantly differ in 5 mM Ca^2+ 48^. This suggests that the RyR isoform difference that we observe is due to regions outside CaMBD2, and further stresses the importance of using full-length RyR in CaM-RyR studies.

CaM association kinetics does not differ between normal CaM and CaM_1234_ for binding to RyR2 at the same nM Ca^2+^. This supports our trilateration results in that CaM is likely in an apo-state when binding to RyR2 in 30 nM Ca^2+^.

By monitoring CaM structural transitions in response to rapid changes between nanomolar and micromolar [Ca^2+^], we determine temporal correlations with CaM-RyR functional modulation in skeletal and cardiac muscle during EC coupling and relaxation.

With both 30 nM to µM and 30 µM to nM Ca^2+^, we observed A-CaM shifts on RyR1 substantially (>50-fold) faster than the observed for A-CaM binding to RyR1 in nM and µM Ca^2+^. This suggests that the FRET shift over time can be attributed to a structural transition while CaM is bound to RyR1, rather than to changes in A-CaM binding to RyR1. Note that the FRET shift was 3.4±0.1 times faster for the [Ca^2+^] shift of nM to µM relative to that of µM to nM. Extrapolating to physiology, this suggests that during EC coupling CaM shifts rapidly to becoming an RyR1 inhibitor, faster than its transition to becoming an RyR1 activator during relaxation.

In contrast, the Ca^2+^ driven structural shift in A-CaM bound to RyR2 appears to be less straightforward. The A-CaM with RyR2 FRET changes driven by rapid shifts in [Ca^2+^] are >10 times faster than the association kinetics, which is consistent with changes in CaM’s structural state, rather than in CaM-RyR2 association. For A-CaM bound to RyR2, the high to low Ca^2+^-driven structural transition, is 4 times slower relative to A-CaM bound to RyR1. This emphasizes that RyR2, relative to RyR1, promotes a Ca-bound state of CaM.

Interestingly, with the 30 nM to µM Ca^2+^ transition, A-CaM bound to RyR2 undergoes a more complex, multiphase shift in FRET, suggesting intermediate structural steps. Most relevant is stage 2 during the 10-20 ms time frame, which is only evident with RyR2, not RyR1. The functional role of stage 2 should become more apparent upon further testing with conditions that shift CaM-RyR2 binding, such as pathologic states of RyR2/CaM, as to be pursued with futures studies.

## METHODS

### Expression, purification and fluorescent labeling of FKBP and CaM

Human FKBP12.6 cDNA was modified by site directed mutagenesis (QuikChange Lightning kit; Agilent Technologies) to introduce one of eight single cysteine mutants (G1C, T14C, E32C, K44C, R49C, Q65C, T85C) into a null cysteine background (C22A and C76I FKBP12.6), FKBP were expressed in *Escherichia coli* BL21(DE3)pLysS, purified, and labeled with AlexaFluor488 (ThermoFisher) as described previously ^35^. Single-cysteine CaM with substitutions within either N lobe (T26C, T34C) or C lobe (Y99C and T110C) were expressed, purified and labeled with AF568 C5 maleimide as described previously^24^. The Ca^2+^-insensitive CaM (CaM_1234_; E-to-A substitutions at 31 67, 104 and 140) cDNA in pET-7 vector was modified by site directed mutagenesis (QuikChange Lightning kit; Agilent Technologies) to introduce one of four single cysteine mutants (T26C, T34C, Y99C and T110C). The CaM_1234_ variants were expressed, purified and labeled with AF568 using the method used to purify FKBP12.6^35^. Stoichiometric labeling of fluorescent CaM and FKBP to ≥95% was demonstrated by ratio of the absorbance of the bound dye relative to SDS-PAGE densitometry. A lack of unlabeled protein was confirmed by MALDI-TOF mass spectrometry.

### SR vesicle isolation

SR membrane vesicles were isolated from pig *longissimus dorsi* and pig ventricular tissue by differential centrifugation of homogenized muscle ^27^. ‘Heavy’ SR, rich in RyR, were isolated by fractionation of skeletal ‘crude’ SR vesicles using a discontinuous sucrose gradient ^27^. All vesicles were stored frozen at -80ºC. SR vesicles were stripped of endogenous CaM by incubation with myosin light chain kinase-derived CaM binding peptide, followed by sedimentation in accordance with ^49^ prior to radioligand and fluorescence binding assays.

### FKBP12.6 binding studies

The binding of F-FKBP to SR membranes was measured as described previously ^35^, with exceptions that SR was incubated with 1-60nM F-FKBP, non-specific FRET was measured by excess addition of 5 µM unlabeled WT FKBP12.6, and AF488 labeled FKBP was used. The binding of FKBPs to SR membranes (0.4 mg/ml) was measured following 90-min incubations in 37 °C buffer containing 150 mM KCl, 20 mM K-PIPES (pH 7.0), 5 mM GSH, 0.1 mg/ml bovine serum albumin, 1 mM EGTA, 65 µM CaCl_2_ (30 nM free Ca^2+^, calculated using Maxchelator), and 1-60 nM FKBP12.6 labeled at indicated sites. For determinations of nonspecific binding, 5 µM unlabeled FKBP12.6 was added to the binding buffer. Bound and free F-FKBP were separated by centrifugation at 100,000 x g. Pellets were resuspended in 5% SDS, 50 mM NaCl, 20 mM Na-PIPES (pH 7.0), and 1 mM EGTA. Bound AF488-FKBP12 was determined from the integrated fluorescence intensity from 510 to 610nm, acquired using a Gemini EM microplate fluorometer (Molecular Devices, Sunnyvale, CA) with excitation at 488 nm and a 495-nm emission long pass filter.

### [^3^H]ryanodine binding

Skeletal and cardiac SR membranes (1 and 3 mg/ml, respectively) were pre-incubated with the indicated range of [AF568-CaM] for 30 min at 4 °C in a solution containing 150mM KCl, 5 mM GSH, 1 µg/ml aprotinin/leupeptin, 1 mM DTT, 1mM EGTA, 65 µM or 1.02 mM CaCl_2_ (30 nM or 30 µM free Ca^2+^, respectively, as determined by MaxChelator), 0.1 mg/ml of BSA, and 20 mM K-PIPES (pH 7.0). The assay solution at 30 nM Ca^2+^ additionally contained 5 mM Na_2_ATP and 5 mM caffeine. Binding of [3H]ryanodine (10 or 15 nM) was determined following a 3-h incubation at 37°C, and filtration through grade GF/B glass microfiber filters (Brandel Inc., Gaithersburg, MD) using a 96-cell Brandel Harvester. In 4 mL of Ecolite scintillation mixture (MP Biomedicals, Solon, OH), the [3H] retained on the filter was measured using a Beckman LS6000 scintillation counter.

### FRET measurements

Skeletal or cardiac SR (0.4mg/ml) membranes were pre-incubated with the donor, 60 nM Alexa Fluor 488-FKBP (D-FKBP), for 90 min at 37ºC in a solution containing 150mM KCl, 5mM GSH, 0.1mg/mL BSA, 1µg/Ml Aprotinin/Leupeptin 1mM DTT and 20mM PIPES (pH 7.0). To remove unbound D-FKBP, the SR membranes were spun at 109,760xg. For trilateration and Ca-driven kinetics studies, D-FKBP treated SR membranes (1mg/mL skeletal SR and 5mg/mL cardiac SR) were incubated for 60 min at 22ºC with 0.8µM Alexa Fluor 568-CaM (A-CaM or A-CaM_1234_) in a *FRET buffer* containing 150mM KCl, 5mM GSH, 1 µg/mL Aprotinin/Leupeptin, 1mM EGTA, 2mM DTT, 65µM CaCl_2_ (30nM free Ca^2+^, as determined by MaxChelator) or 1.02mM CaCl_2_ (30µM free Ca, as determined by MaxChelator) and 0.1mg/mL of BSA and 20mM K-PIPES (pH 7.0). For CaM association studies using a stopped-flow instrument^44, 45^, one syringe was loaded with loaded with D-FKBP treated SR (2mg/mL skeletal SR and 6mg/mL cardiac SR), with the second syringe loaded with 1.6 µM A-CaM, both with *FRET buffer* for either 30 nM or µM free Ca^2+^. For Ca-driven CaM-RyR structural transition studies, one syringe was loaded with D-FKBP treated SR (2mg/mL skeletal SR and 6mg/mL cardiac SR) and 1.6 µM A-CaM in *FRET buffer*, with the second syringe loaded with Ca^2+^ or EGTA in *FRET buffer* solution to shift the final free Ca^2+^ from 30 nM to µM Ca^2+^ or 30 µM to nM Ca^2+^ (as determined by MaxChelator).

For data used in trilateration, we acquired fluorescence decays using time-correlated single-photon counting, as described previously^50^. For kinetics studies, we acquired fluorescence decays using direct waveform recording, as described previously^44, 45^. The instrument response function (IRF) was acquired using water in the cuvette. Global multi-exponential analysis of the FLT-FRET data was used to test a series of structural models, as extensively described previously^50, 51^.

### Simulation of donor probe positions

To interpret the FKBP-CaM FRET data, we determined the location of the donor AF488 probes when attached to FKBP in RyR1 and RyR2. This was done similarly to what we had previously done^38^ except that here we used the higher-resolution RyR structures currently available. For the computational studies of RyR1, the structure chosen was the 3.6-Å closed state (PDB ID: 5T15) and the corresponding cryo-EM density map (EMDB ID: 8342)^1^. For RyR2, we used the 3.6-Å structure corresponding to the closed state (PDB ID: 6IJ8) and the corresponding cryo-EM density map (EMDB ID: 9833)^4^.

For molecular simulations of the donor probes, we need to take into account the protein volume corresponding to the region around FKBP. Near FKBP is the DR2 region, which, due to disorder, has not been resolved in any cryo-EM structures for RyR. To allow for that region, we used the protein density of the cryo-EM maps represented as dummy atoms in the simulations. The grid points of the new map were converted to dummy atoms using the Situs software package “vol2pdb” program^52^ and saved in a pdb file. Grid-points overlapping with the coordinates of FKBP were removed. To speed up calculations, only dummy atoms within 30 Å from FKBP were used to restrict the conformational space of the fluorescent probe, and all grid-points that are completely buried were deleted.

In the atomic structure of FKBP12.6, the eight residues at position 1, 6, 14, 32, 44, 49, 65, and 85 that were used for probe attachment in the fluorescence experiments were mutated to cysteines using Discovery Studio Visualizer 2019 (BIOVIA, Dassault Systèmes, San Diego, CA). AF488 was attached, respectively, to each of the eight cysteines, generating eight starting conformations for simulated annealing using the software package CNS 1.3^53^. The simulated annealing protocol has been described in detail and validated previously^38^. With the assumption that the fluorescence probe undergoes fast dynamics in relation to the fluorescence lifetime^54^, means that we can use the average position the chromophore’s center, as calculated from simulated annealing.

### Calculation of predicted acceptor probe locations

To calculate the expected distances between donor and acceptor probes in the cryo-EM structures we developed a new fast method to determine the predicted location of the acceptor probes on CaM. As this information is just used for comparison with experimental FRET distance it would not be necessary to run the computationally costly simulated annealing calculations to determine the probe locations. Data from the previous probe simulations using simulated annealing of AF488 attached to FKBP were analyzed for the distance between the center of the chromophore back to the Cβ atom of the Cys which the probe is attached to. These distances were binned to generate a probability distribution. The probe center to the Cβ atom distance was found to be between 8 and 20 Å with an average value of ∼15 Å. A region in space that the probe can sample was determined by selecting points that did not clash with atoms of the protein around the attachment site of the probe, from points on concentric spheres of evenly spaced points. A radius of 6 Å for the probe was used. These selected points in space were combined with the probability distribution previously determined to calculate the average probe location. The method was validated by calculating average probe positions for AF488 on FKBP and was found to be quite accurate as it could predict probe locations that were on average within 2-3 Å from the locations determined by simulated annealing.

### Trilateration

From the simulated-annealing calculations above, we obtained coordinates for the probe locations, which we used as effective AF488 chromophore locations for trilateration. We used these coordinates for each of the eight FKBP attachment sites (1, 6, 14, 32, 44, 49, 65, and 85), in combination with distances calculated from FRET, to determine a locus in space corresponding to the acceptor attached to each of the four labeled sites on CaM (26, 34, 99, and 110). We have developed software that performs trilateration using an arbitrary number of donor sites and distances. Our new trilateration approach takes into account Gaussian distance distributions as fitted to the FLT-FRET measurement for each donor position. The normalized sum of the distance distributions from all of the donor sites to one acceptor is generated as a probability map describing the acceptor’s location. This volumetric map is saved in MRC/CCP4-format, which can be displayed using common molecular graphics software. The trilateration software can be obtained from the authors. To focus the results to just the region near RyR the probability map was masked to the region within 20 Å from the RyR surface using UCSF Chimera 1.16^55^. Chimera was also used to calculate the volume and center positions of the trilaterated loci and determine the iso value setting for displaying the loci at the selected 1000 Å^3^ volume. Molecular structure figures were created using VMD 1.9.3^56^.

#### Statistics

Sample means are from three or more independent experiments; numbers of observations (N) are given in the figure legends. Each experiment was repeated using at least two independent SR membrane preparations, isolated from different animals. Average data are provided as mean ± SE. Statistical significance was evaluated by use of either paired or unpaired Student’s t-test or analysis of variance as appropriate.

