## Supplementary Information for "Kinetics and Mapping of Ca-driven Calmodulin conformations on Skeletal and Cardiac Muscle Ryanodine Receptors"

From the [1]Department of Biochemistry, Molecular Biology & Biophysics, University of Minnesota, Minneapolis, MN, USA 55455;
[2]Department of Pharmacology, University of California at Davis, Davis, CA, USA 95616.
[#]Both authors contributed equally to this work.

**Supplementary Table 1. Comparison between cryo-EM and FRET-determined distances (used for trilateration) between probe sites attached to indicated (X) site on FKBP and N-lobe residues (T26 and T34) on CaM, both bound to RyR1 in skeletal SR membranes.** Cryo-EM PDB 6X32 and 7TZC were used to modeled probe sites on FKBP and CaM, both bound to purified RyR1, and then measure distances between probes sites. FLT-FRET was used to model Gaussian distributions between dye probe sites, and the values in the table indicate the values used for trilateration.

**RyR1 FKBP-X to CaM-26**

| FKBP | Apo-CaM (closed) 6X32 Dist. | Ca-CaM (closed) 7TZC Dist. | CaM nM $Ca^{2+}$ FRET Dist. | FWHM | CaM μM $Ca^{2+}$ FRET Dist. | FWHM | $CaM_{1234}$ nM $Ca^{2+}$ FRET Dist. | FWHM | $CaM_{1234}$ μM $Ca^{2+}$ FRET Dist. | FWHM |
|---|---|---|---|---|---|---|---|---|---|---|
| 1 | 39.0 | 35.8 | 52.8 | 19.8 | 75.0 | 17.5 | 52.0 | 15.2 | 52.0 | 15.7 |
| 6 | 55.0 | 51.3 | 57.7 | 20.8 | 59.9 | 31.5 | 58.2 | 27.0 | 71.6 | 17.8 |
| 14 | 49.5 | 46.3 | 56.4 | 12.6 | 57.0 | 18.9 | 68.5 | 18.2 | 62.8 | 26.9 |
| 32 | 42.8 | 37.6 | 45.6 | 29.6 | 60.8 | 48.0 | 44.6 | 29.6 | 45.7 | 26.8 |
| 44 | 42.5 | 37.3 | 54.6 | 18.8 | 52.6 | 21.1 | 56.3 | 21.9 | 57.1 | 23.5 |
| 49 | 28.1 | 24.3 | 49.3 | 14.4 | 48.6 | 20.8 | 51.3 | 19.1 | 51.4 | 19.9 |
| 65 | 45.8 | 42.7 | 55.4 | 17.8 | 63.5 | 27.8 | 73.6 | 25.9 | 52.0 | 11.1 |
| 85 | 26.7 | 22.5 | 38.4 | 31.7 | 46.7 | 57.9 | 39.2 | 28.0 | 40.6 | 18.5 |

**RyR1 FKBP-X to CaM-34**

| FKBP | Apo-CaM (closed) 6X32 Dist. | Ca-CaM (closed) 7TZC Dist. | CaM nM $Ca^{2+}$ FRET Dist. | FWHM | CaM μM $Ca^{2+}$ FRET Dist. | FWHM | $CaM_{1234}$ nM $Ca^{2+}$ FRET Dist. | FWHM | $CaM_{1234}$ μM $Ca^{2+}$ FRET Dist. | FWHM |
|---|---|---|---|---|---|---|---|---|---|---|
| 1 | 52.2 | 43.0 | 68.4 | 24.0 | 72.6 | 18.3 | 66.3 | 29.6 | 60.2 | 20.9 |
| 6 | 64.4 | 55.7 | 61.2 | 19.3 | 61.1 | 20.3 | 58.9 | 22.2 | 55.7 | 10.4 |
| 14 | 52.4 | 45.4 | 52.5 | 15.7 | 54.0 | 15.7 | 51.1 | 10.6 | 52.4 | 15.4 |
| 32 | 62.2 | 52.5 | 61.9 | 50.2 | 67.8 | 50.2 | 58.9 | 18.1 | 59.5 | 16.2 |
| 44 | 49.0 | 41.6 | 52.3 | 18.7 | 50.8 | 18.7 | 55.3 | 23.8 | 51.8 | 12.9 |
| 49 | 33.1 | 25.2 | 48.0 | 19.3 | 46.0 | 19.3 | 46.4 | 19.7 | 45.2 | 21.8 |
| 65 | 55.7 | 47.0 | 69.0 | 22.0 | 61.3 | 20.1 | 57.2 | 15.0 | 55.4 | 8.9 |
| 85 | 46.2 | 36.8 | 53.9 | 24.9 | 52.3 | 27.2 | 49.3 | 28.0 | 49.5 | 25.0 |

**Supplementary Table 2. Comparison between cryo-EM and FRET-determined distances (used for trilateration) between probe sites attached to indicated (X) site on FKBP and C-lobe residues (T26 and T34) on CaM, both bound to RyR1 in skeletal SR membranes.** Cryo-EM PDB 6X32 and 7TZC were used to modeled probe sites on FKBP and CaM, both bound to purified RyR1, and then measure distances between probes sites. FLT-FRET was used to model Gaussian distributions between dye probe sites, and the values in the table indicate the values used for trilateration.

**RyR1 FKBP-X to CaM-99**

| FKBP | Apo-CaM (closed) 6X32 Dist. | Ca-CaM (closed) 7TZC Dist. | CaM nM $Ca^{2+}$ FRET Dist. | FWHM | CaM μM $Ca^{2+}$ FRET Dist. | FWHM | $CaM_{1234}$ nM $Ca^{2+}$ FRET Dist. | FWHM | $CaM_{1234}$ μM $Ca^{2+}$ FRET Dist. | FWHM |
|---|---|---|---|---|---|---|---|---|---|---|
| 1 | 100.6 | 96.6 | 101.1 | 27.1 | 95.7 | 20.9 | 102.8 | 31.3 | 98.4 | 35.9 |
| 6 | 110.4 | 107.0 | 101.3 | 21.3 | 98.8 | 27.5 | 111.1 | 32.6 | 101.3 | 28.0 |
| 14 | 92.9 | 90.2 | 87.7 | 14.3 | 86.9 | 18.2 | 92.9 | 21.0 | 93.4 | 14.3 |
| 32 | 102.6 | 98.0 | 90.4 | 12.9 | 88.6 | 20.2 | 92.5 | 23.4 | 89.6 | 16.1 |
| 44 | 78.6 | 75.4 | 86.9 | 9.7 | 78.9 | 12.3 | 86.5 | 12.9 | 87.0 | 21.3 |
| 49 | 71.5 | 68.1 | 83.7 | 11.5 | 76.9 | 14.4 | 86.3 | 18.7 | 84.7 | 17.9 |
| 65 | 103.5 | 99.9 | 103.5 | 20.5 | 94.2 | 17.7 | 106.1 | 28.3 | 103.5 | 29.8 |
| 85 | 89.2 | 84.7 | 81.7 | 10.4 | 81.9 | 19.8 | 88.7 | 23.2 | 89.5 | 25.4 |

**RyR1 FKBP-X to CaM-110**

| FKBP | Apo-CaM (closed) 6X32 Dist. | Ca-CaM (closed) 7TZC Dist. | CaM nM $Ca^{2+}$ FRET Dist. | FWHM | CaM μM $Ca^{2+}$ FRET Dist. | FWHM | $CaM_{1234}$ nM $Ca^{2+}$ FRET Dist. | FWHM | $CaM_{1234}$ μM $Ca^{2+}$ FRET Dist. | FWHM |
|---|---|---|---|---|---|---|---|---|---|---|
| 1 | 106.3 | 103.4 | 108.9 | 35.6 | 104.9 | 42.9 | 106.6 | 33.9 | 103.6 | 26.9 |
| 6 | 115.3 | 111.8 | 112.5 | 25.5 | 96.0 | 26.4 | 112.6 | 23.0 | 106.0 | 20.6 |
| 14 | 99.2 | 95.8 | 88.0 | 13.0 | 80.8 | 10.6 | 97.2 | 20.5 | 95.2 | 20.0 |
| 32 | 101.0 | 97.7 | 96.6 | 10.6 | 81.3 | 13.2 | 100.4 | 21.2 | 101.2 | 24.7 |
| 44 | 77.4 | 73.3 | 82.1 | 10.0 | 74.0 | 14.4 | 87.4 | 11.6 | 87.1 | 20.4 |
| 49 | 76.2 | 72.9 | 86.8 | 14.5 | 74.2 | 14.2 | 92.9 | 22.8 | 91.5 | 18.5 |
| 65 | 109.7 | 106.6 | 106.8 | 20.6 | 90.9 | 17.5 | 111.9 | 28.8 | 108.4 | 28.5 |
| 85 | 99.4 | 88.7 | 87.8 | 17.4 | 81.1 | 29.0 | 95.7 | 19.4 | 94.6 | 26.2 |

**Supplementary Table 3. Comparison between cryo-EM and FRET-determined distances (used for trilateration) between probe sites attached to indicated (X) site on FKBP and N-lobe residues (T26 and T34) on CaM, both bound to RyR2 in cardiac SR membranes.** Cryo-EM PDB 6JI8 and 6JV2 were used to modeled probe sites on FKBP and CaM, both bound to purified RyR1, and then measure distances between probes sites. FLT-FRET was used to model Gaussian distributions between dye probe sites, and the values in the table indicate the values used for trilateration.

**RyR2 FKBP-X to CaM-26**

| | Apo-CaM (closed) 6JI8 | Ca-CaM (closed) 6JV2 | CaM nM $Ca^{2+}$ | | CaM μM $Ca^{2+}$ | | $CaM_{1234}$ nM $Ca^{2+}$ | | $CaM_{1234}$ μM $Ca^{2+}$ | |
|---|---|---|---|---|---|---|---|---|---|---|
| FKBP | Dist. | Dist. | FRET Dist. | FWHM | FRET Dist. | FWHM | FRET Dist. | FWHM | FRET Dist. | FWHM |
| 1 | 46.0 | 76.1 | 52.0 | 9.7 | 57.9 | 13.2 | 52.4 | 12.0 | 51.7 | 9.3 |
| 6 | 58.5 | 84.6 | 55.3 | 10.5 | 54.6 | 19.9 | 60.6 | 15.2 | 60.3 | 15.7 |
| 14 | 56.1 | 74.2 | 54.0 | 10.7 | 52.7 | 35.1 | 68.3 | 19.3 | 63.1 | 29.2 |
| 32 | 44.5 | 73.9 | 57.2 | 42.9 | 57.4 | 48.2 | 44.7 | 32.9 | 45.3 | 21.6 |
| 44 | 43.6 | 57.8 | 38.0 | 36.4 | 42.4 | 31.8 | 52.9 | 21.0 | 52.3 | 21.8 |
| 49 | 31.1 | 48.4 | 45.0 | 20.3 | 50.7 | 22.4 | 48.4 | 17.7 | 44.7 | 26.1 |
| 65 | 52.4 | 79.0 | 59.9 | 15.9 | 61.0 | 18.6 | 61.2 | 16.9 | 61.6 | 15.7 |
| 85 | 27.6 | 57.7 | 32.2 | 35.1 | 50.2 | 36.9 | 36.6 | 34.8 | 41.6 | 22.7 |

**RyR2 FKBP-X to CaM-34**

| | Apo-CaM (closed) 6JI8 | Ca-CaM (closed) 6JV2 | CaM nM $Ca^{2+}$ | | CaM μM $Ca^{2+}$ | | $CaM_{1234}$ nM $Ca^{2+}$ | | $CaM_{1234}$ μM $Ca^{2+}$ | |
|---|---|---|---|---|---|---|---|---|---|---|
| FKBP | Dist. | Dist. | FRET Dist. | FWHM | FRET Dist. | FWHM | FRET Dist. | FWHM | FRET Dist. | FWHM |
| 1 | 58.6 | 95.9 | 63.3 | 9.6 | 59.7 | 12.9 | 56.5 | 13.5 | 60.8 | 9.9 |
| 6 | 66.9 | 106.1 | 62.8 | 10.0 | 60.4 | 16.7 | 63.9 | 13.4 | 63.4 | 19.3 |
| 14 | 57.0 | 96.0 | 54.5 | 31.3 | 58.6 | 32.4 | 50.6 | 14.2 | 52.4 | 18.8 |
| 32 | 62.9 | 90.4 | 104.2 | 51.6 | 74.9 | 33.2 | 62.8 | 22.5 | 64.7 | 25.1 |
| 44 | 47.4 | 75.6 | 51.2 | 26.8 | 52.6 | 24.6 | 52.9 | 22.6 | 49.1 | 15.6 |
| 49 | 32.8 | 70.0 | 50.7 | 28.8 | 46.8 | 40.5 | 44.4 | 17.5 | 39.2 | 28.2 |
| 65 | 61.0 | 100.7 | 62.4 | 7.3 | 62.7 | 18.0 | 62.4 | 9.7 | 62.2 | 9.3 |
| 85 | 46.7 | 75.2 | 51.0 | 15.4 | 51.4 | 39.4 | 48.5 | 26.4 | 51.0 | 20.1 |

**Supplementary Table 4. Comparison between cryo-EM and FRET-determined distances (used for trilateration) between probe sites attached to indicated (X) site on FKBP and C-lobe residues (T26 and T34) on CaM, both bound to RyR2 in cardiac SR membranes.** Cryo-EM PDB 6JI8 and 6JV2 were used to modeled probe sites on FKBP and CaM, both bound to purified RyR1, and then measure distances between probes sites. FLT-FRET was used to model Gaussian distributions between dye probe sites, and the values in the table indicate the values used for trilateration.

**RyR2 FKBP-X to CaM-99**

| | Apo-CaM (closed) | Ca-CaM (closed) | | | | | | | | |
|---|---|---|---|---|---|---|---|---|---|---|
| | 6JI8 | 6JV2 | CaM nM $Ca^{2+}$ | | CaM μM $Ca^{2+}$ | | $CaM_{1234}$ nM $Ca^{2+}$ | | $CaM_{1234}$ μM $Ca^{2+}$ | |
| FKBP | Dist. | Dist. | FRET Dist. | FWHM | FRET Dist. | FWHM | FRET Dist. | FWHM | FRET Dist. | FWHM |
| 1 | 106.3 | 123.5 | 115.8 | 61.6 | 101.1 | 46.4 | 104.9 | 46.7 | 114.8 | 57.5 |
| 6 | 112.9 | 128.3 | 110.8 | 40.8 | 97.9 | 27.5 | 93.1 | 30.8 | 96.5 | 24.8 |
| 14 | 97.3 | 110.3 | 86.0 | 24.2 | 81.6 | 17.3 | 95.1 | 27.4 | 96.1 | 23.1 |
| 32 | 104.2 | 121.9 | 101.8 | 5.0 | 94.2 | 5.0 | 93.4 | 25.3 | 86.3 | 24.6 |
| 44 | 78.2 | 91.4 | 91.0 | 30.6 | 85.9 | 22.6 | 86.2 | 19.7 | 84.3 | 15.3 |
| 49 | 74.0 | 89.1 | 90.6 | 13.0 | 88.8 | 17.8 | 83.9 | 23.5 | 86.7 | 25.1 |
| 65 | 108.2 | 124.1 | 108.6 | 37.3 | 101.6 | 34.8 | 95.8 | 33.3 | 86.2 | 24.5 |
| 85 | 89.4 | 107.7 | 96.2 | 34.8 | 94.9 | 32.4 | 86.2 | 15.8 | 90.1 | 34.2 |

**RyR2 FKBP-X to CaM-110**

| | Apo-CaM (closed) | Ca-CaM (closed) | | | | | | | | |
|---|---|---|---|---|---|---|---|---|---|---|
| | 6JI8 | 6JV2 | CaM nM $Ca^{2+}$ | | CaM μM $Ca^{2+}$ | | $CaM_{1234}$ nM $Ca^{2+}$ | | $CaM_{1234}$ μM $Ca^{2+}$ | |
| FKBP | Dist. | Dist. | FRET Dist. | FWHM | FRET Dist. | FWHM | FRET Dist. | FWHM | FRET Dist. | FWHM |
| 1 | 106.2 | 114.0 | 119.6 | 62.8 | 104.2 | 51.7 | 119.8 | 62.1 | 116.5 | 61.3 |
| 6 | 112.7 | 121.1 | 110.8 | 23.2 | 94.8 | 24.6 | 91.9 | 20.8 | 98.4 | 25.7 |
| 14 | 99.1 | 107.0 | 92.9 | 34.5 | 84.2 | 24.8 | 96.7 | 24.1 | 98.1 | 22.2 |
| 32 | 98.4 | 107.5 | 107.6 | 5.0 | 89.3 | 35.0 | 100.5 | 37.2 | 101.8 | 37.5 |
| 44 | 74.2 | 83.6 | 87.8 | 30.9 | 77.9 | 21.8 | 85.0 | 18.9 | 87.6 | 21.6 |
| 49 | 76.3 | 83.5 | 94.4 | 5.7 | 84.1 | 17.2 | 89.0 | 21.9 | 94.9 | 20.2 |
| 65 | 109.0 | 116.9 | 111.8 | 36.7 | 98.6 | 32.0 | 94.4 | 26.1 | 94.9 | 35.4 |
| 85 | 86.7 | 94.5 | 100.5 | 31.6 | 92.3 | 39.0 | 89.3 | 13.9 | 85.2 | 18.5 |

**Supplementary Table 5. Comparison between cryo-EM and trilateration Loci (from FRET data) distances between probe sites attached to indicated (X) site on FKBP and N-lobe residues (T26 and T34) on CaM, both bound to RyR1 in skeletal SR membranes.** Cryo-EM PDB 6X32 and 7TZC were used to modeled probe sites on FKBP and CaM, both bound to purified RyR1, and then measure distances between probes sites.

**RyR1 FKBP-X to CaM-26**

| FKBP | Apo-CaM (closed) 6X32 Dist. | Ca-CaM (closed) 7TZC Dist. | CaM nM $Ca^{2+}$ Locus Dist. | CaM µ$Ca^{2+}$ Locus Dist. | $CaM_{1234}$ nM $Ca^{2+}$ Locus Dist. | $CaM_{1234}$ µM $Ca^{2+}$ Locus Dist. |
|---|---|---|---|---|---|---|
| 1 | 39.0 | 35.8 | 47.3 | 65.1 | 52.8 | 49.1 |
| 6 | 55.0 | 51.3 | 64.2 | 76.6 | 71.5 | 67.2 |
| 14 | 49.5 | 46.3 | 58.2 | 62.6 | 67.8 | 62.6 |
| 32 | 42.8 | 37.6 | 53.2 | 68.0 | 51.7 | 53.7 |
| 44 | 42.5 | 37.3 | 53.6 | 49.9 | 59.0 | 58.0 |
| 49 | 28.1 | 24.3 | 38.1 | 39.1 | 45.9 | 42.7 |
| 65 | 45.8 | 42.7 | 54.4 | 68.9 | 61.8 | 57.1 |
| 85 | 26.7 | 22.5 | 36.5 | 53.9 | 36.5 | 37.2 |

**RyR1 FKBP-X to CaM-34**

| FKBP | Apo-CaM (closed) 6X32 Dist. | Ca-CaM (closed) 7TZC Dist. | CaM nM $Ca^{2+}$ Locus Dist. | CaM µ$Ca^{2+}$ Locus Dist. | $CaM_{1234}$ nM $Ca^{2+}$ Locus Dist. | $CaM_{1234}$ µM $Ca^{2+}$ Locus Dist. |
|---|---|---|---|---|---|---|
| 1 | 52.2 | 43.0 | 59.8 | 56.7 | 52.3 | 50.5 |
| 6 | 64.4 | 55.7 | 71.4 | 69.1 | 64.4 | 64.6 |
| 14 | 52.4 | 45.4 | 57.8 | 56.5 | 51.9 | 54.8 |
| 32 | 62.2 | 52.5 | 66.5 | 64.0 | 59.6 | 58.4 |
| 44 | 49.0 | 41.6 | 49.7 | 49.6 | 44.7 | 50.0 |
| 49 | 33.1 | 25.2 | 36.1 | 35.2 | 30.1 | 34.3 |
| 65 | 55.7 | 47.0 | 63.3 | 60.6 | 56.1 | 55.5 |
| 85 | 46.2 | 36.8 | 51.2 | 48.4 | 44.3 | 42.1 |

**Supplementary Table 6. Comparison between cryo-EM and trilateration Loci (from FRET data) distances between probe sites attached to indicated (X) site on FKBP and N-lobe residues (T99 and T110) on CaM, both bound to RyR1 in skeletal SR membranes.** Cryo-EM PDB 6X32 and 7TZC were used to modeled probe sites on FKBP and CaM, both bound to purified RyR1, and then measure distances between probes sites.

**RyR1 FKBP-X to CaM-99**

| FKBP | Apo-CaM (closed) 6X32 Dist. | Ca-CaM (closed) 7TZC Dist. | CaM nM $Ca^{2+}$ Locus Dist. | CaM $\mu Ca^{2+}$ Locus Dist. | $CaM_{1234}$ nM $Ca^{2+}$ Locus Dist. | $CaM_{1234}$ $\mu M$ $Ca^{2+}$ Locus Dist. |
|---|---|---|---|---|---|---|
| 1 | 100.6 | 96.6 | 96.5 | 93.3 | 102.7 | 95.3 |
| 6 | 110.4 | 107.0 | 108.6 | 106.8 | 116.1 | 108.8 |
| 14 | 92.9 | 90.2 | 94.8 | 93.5 | 103.0 | 95.9 |
| 32 | 102.6 | 98.0 | 92.0 | 91.4 | 98.4 | 92.1 |
| 44 | 78.6 | 75.4 | 75.5 | 77.7 | 85.0 | 79.0 |
| 49 | 71.5 | 68.1 | 70.3 | 69.4 | 78.4 | 71.4 |
| 65 | 103.5 | 99.9 | 101.6 | 98.8 | 108.5 | 101.0 |
| 85 | 89.2 | 84.7 | 80.6 | 78.2 | 86.3 | 79.4 |

**RyR1 FKBP-X to CaM-110**

| FKBP | Apo-CaM (closed) 6X32 Dist. | Ca-CaM (closed) 7TZC Dist. | CaM nM $Ca^{2+}$ Locus Dist. | CaM $\mu Ca^{2+}$ Locus Dist. | $CaM_{1234}$ nM $Ca^{2+}$ Locus Dist. | $CaM_{1234}$ $\mu M$ $Ca^{2+}$ Locus Dist. |
|---|---|---|---|---|---|---|
| 1 | 106.3 | 103.4 | 101.8 | 83.6 | 106.8 | 103.0 |
| 6 | 115.3 | 111.8 | 114.4 | 96.4 | 118.7 | 114.1 |
| 14 | 99.2 | 95.8 | 100.8 | 82.7 | 103.8 | 98.1 |
| 32 | 101.0 | 97.7 | 96.9 | 84.3 | 103.9 | 102.4 |
| 44 | 77.4 | 73.3 | 81.6 | 69.0 | 86.1 | 81.8 |
| 49 | 76.2 | 72.9 | 76.2 | 59.2 | 80.0 | 75.2 |
| 65 | 109.7 | 106.6 | 107.2 | 88.4 | 111.5 | 107.0 |
| 85 | 99.4 | 88.7 | 85.5 | 70.3 | 91.7 | 89.6 |

**Supplementary Table 7. Comparison between cryo-EM and trilateration Loci (from FRET data) distances between probe sites attached to indicated (X) site on FKBP and N-lobe residues (T26 and T34) on CaM, both bound to RyR1 in skeletal SR membranes.** Cryo-EM PDB 6JI8 and 6JV2 were used to modeled probe sites on FKBP and CaM, both bound to purified RyR1, and then measure distances between probes sites.

**RyR2 FKBP-X to CaM-26**

| FKBP | Apo-CaM (closed) 6JI8 Dist. | Ca-CaM (closed) 6JV2 Dist. | CaM nM $Ca^{2+}$ Locus Dist. | CaM $\mu Ca^{2+}$ Locus Dist. | $CaM_{1234}$ nM $Ca^{2+}$ Locus Dist. | $CaM_{1234}$ $\mu M$ $Ca^{2+}$ Locus Dist. |
|---|---|---|---|---|---|---|
| 1 | 46.0 | 76.1 | 50.5 | 57.0 | 52.3 | 51.3 |
| 6 | 58.5 | 84.6 | 60.0 | 64.2 | 66.1 | 64.4 |
| 14 | 56.1 | 74.2 | 52.8 | 52.8 | 64.6 | 62.1 |
| 32 | 44.5 | 73.9 | 51.3 | 60.4 | 51.3 | 51.2 |
| 44 | 43.6 | 57.8 | 39.5 | 40.6 | 53.8 | 51.5 |
| 49 | 31.1 | 48.4 | 26.7 | 27.7 | 40.2 | 37.5 |
| 65 | 52.4 | 79.0 | 54.3 | 58.8 | 59.5 | 57.9 |
| 85 | 27.6 | 57.7 | 35.4 | 45.1 | 34.0 | 33.8 |

**RyR2 FKBP-X to CaM-34**

| FKBP | Apo-CaM (closed) 6JI8 Dist. | Ca-CaM (closed) 6JV2 Dist. | CaM nM $Ca^{2+}$ Locus Dist. | CaM $\mu Ca^{2+}$ Locus Dist. | $CaM_{1234}$ nM $Ca^{2+}$ Locus Dist. | $CaM_{1234}$ $\mu M$ $Ca^{2+}$ Locus Dist. |
|---|---|---|---|---|---|---|
| 1 | 58.6 | 95.9 | 60.8 | 58.7 | 57.2 | 60.0 |
| 6 | 66.9 | 106.1 | 66.4 | 67.6 | 66.3 | 67.7 |
| 14 | 57.0 | 96.0 | 52.7 | 58.5 | 57.8 | 56.8 |
| 32 | 62.9 | 90.4 | 63.6 | 63.1 | 61.6 | 63.9 |
| 44 | 47.4 | 75.6 | 38.4 | 49.4 | 48.9 | 46.0 |
| 49 | 32.8 | 70.0 | 28.1 | 34.4 | 33.7 | 32.3 |
| 65 | 61.0 | 100.7 | 61.5 | 61.5 | 60.2 | 62.0 |
| 85 | 46.7 | 75.2 | 49.2 | 46.7 | 45.1 | 48.0 |

**Supplementary Table 8. Comparison between cryo-EM and trilateration Loci (from FRET data) distances between probe sites attached to indicated (X) site on FKBP and N-lobe residues (T99 and T110) on CaM, both bound to RyR1 in skeletal SR membranes.** Cryo-EM PDB 6JI8 and 6JV2 were used to modeled probe sites on FKBP and CaM, both bound to purified RyR1, and then measure distances between probes sites.

**RyR2 FKBP-X to CaM-99**

| FKBP | Apo-CaM (closed) 6JI8 Dist. | Ca-CaM (closed) 6JV2 Dist. | CaM nM $Ca^{2+}$ Locus Dist. | CaM μ$Ca^{2+}$ Locus Dist. | $CaM_{1234}$ nM $Ca^{2+}$ Locus Dist. | $CaM_{1234}$ μM $Ca^{2+}$ Locus Dist. |
|---|---|---|---|---|---|---|
| 1 | 106.3 | 123.5 | 104.2 | 94.6 | 104.5 | 101.1 |
| 6 | 112.9 | 128.3 | 111.0 | 99.7 | 113.9 | 111.1 |
| 14 | 97.3 | 110.3 | 96.1 | 83.1 | 102.3 | 100.2 |
| 32 | 104.2 | 121.9 | 100.7 | 93.5 | 99.6 | 96.3 |
| 44 | 78.2 | 91.4 | 75.3 | 64.0 | 81.9 | 80.4 |
| 49 | 74.0 | 89.1 | 72.4 | 60.6 | 76.8 | 74.5 |
| 65 | 108.2 | 124.1 | 106.5 | 95.4 | 108.7 | 105.7 |
| 85 | 89.4 | 107.7 | 86.4 | 79.4 | 84.5 | 81.0 |

**RyR2 FKBP-X to CaM-110**

| FKBP | Apo-CaM (closed) 6JI8 Dist. | Ca-CaM (closed) 6JV2 Dist. | CaM nM $Ca^{2+}$ Locus Dist. | CaM μ$Ca^{2+}$ Locus Dist. | $CaM_{1234}$ nM $Ca^{2+}$ Locus Dist. | $CaM_{1234}$ μM $Ca^{2+}$ Locus Dist. |
|---|---|---|---|---|---|---|
| 1 | 106.2 | 114.0 | 109.8 | 94.5 | 108.0 | 103.3 |
| 6 | 112.7 | 121.1 | 115.5 | 102.4 | 116.5 | 112.3 |
| 14 | 99.1 | 107.0 | 99.1 | 88.7 | 103.6 | 99.8 |
| 32 | 98.4 | 107.5 | 107.0 | 93.9 | 103.5 | 100.1 |
| 44 | 74.2 | 83.6 | 78.7 | 72.0 | 83.1 | 80.9 |
| 49 | 76.3 | 83.5 | 76.6 | 64.4 | 78.8 | 74.7 |
| 65 | 109.0 | 116.9 | 111.3 | 97.1 | 111.6 | 107.0 |
| 85 | 86.7 | 94.5 | 93.0 | 78.1 | 88.6 | 84.5 |

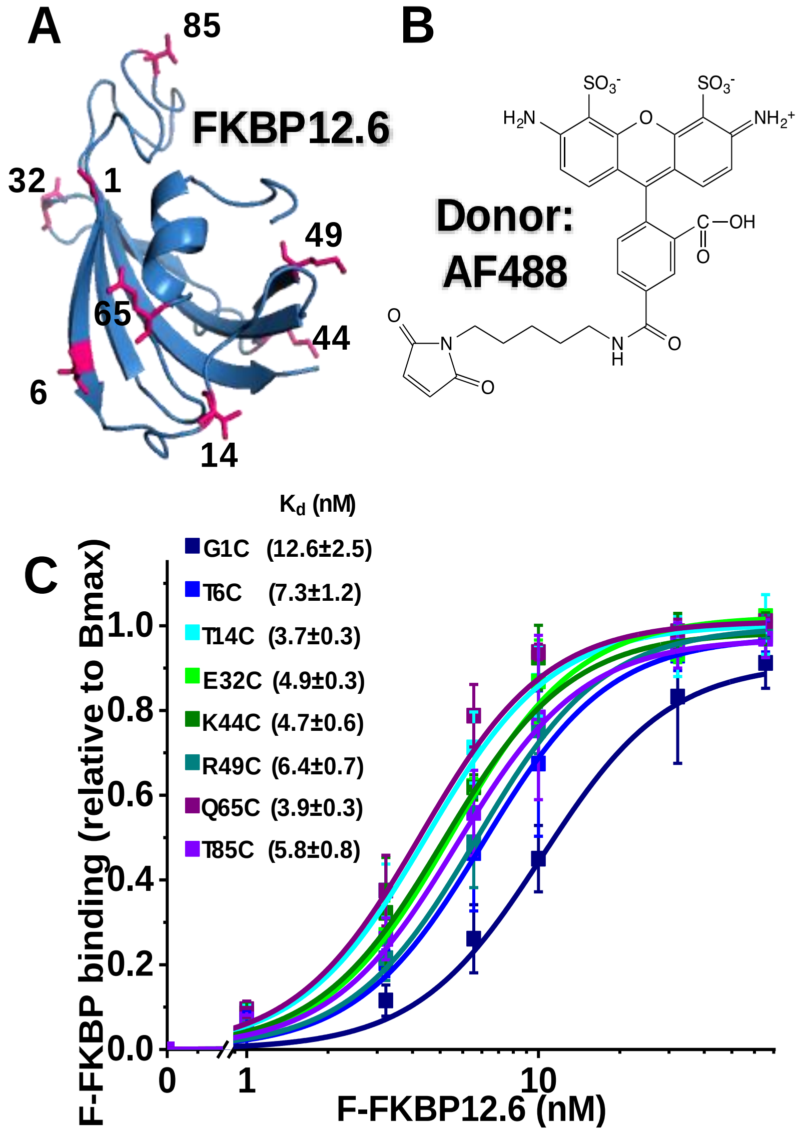


**Supplementary Fig. 1 Eight labeling sites on FKBP12.6 and impact on binding to RyR**. A) Crystal structure of FKBP12.6 with sites mutated to Cys for AF488 C5 maleimide labeling shown in pink. B) chemical structure of AF488. C) Fluorescence intensity indicative of D-FKBP binding to skeletal SR following sedimentation to remove unbound D-FKBP. Data shown as mean ±SE, n = 5 experiments. $K_d$ values from Hill fit to data using Origin 2015 software.

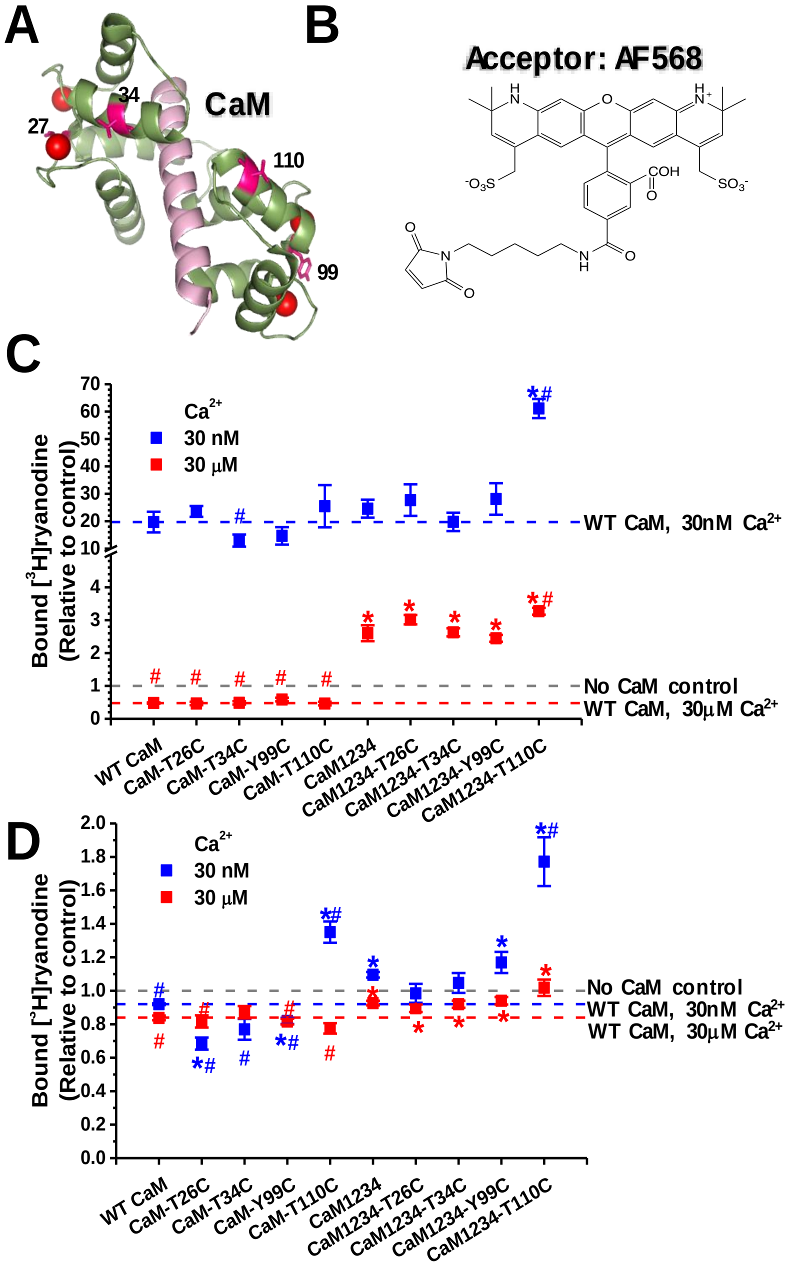


**Supplementary Fig. 2 Four labeling sites on $Ca^{2+}$-sensitive and -insensitive CaM and functional impact of labeling on binding to RyR.** A) Crystal structure of CaM with labeling sites (pink) for shown FRET donor probe AF568. B) Chemical structure of AF568. C and D) SR membranes from skeletal (C) or cardiac (D) muscle were incubated with indicated CaM (0 or 800 nM) at 30 nM (blue) or 30 μM (red) $Ca^{2+}$ in the presence of [$^3$H]ryanodine. The fraction of [$^3$H]ryanodine binding is relative to "no CaM" control indicated with gray line, with the WT CaM effect at 30nM and 30 μM $Ca^{2+}$ indicated with blue and red dotted lines, respectively. Data shown as mean ±SE, n = 4-9 experiments. *Significant differences relative to WT-CaM control, $p<0.05$. #Significant differences relative to Ca-insensitive CaM ($CaM_{1234}$), $p<0.05$.

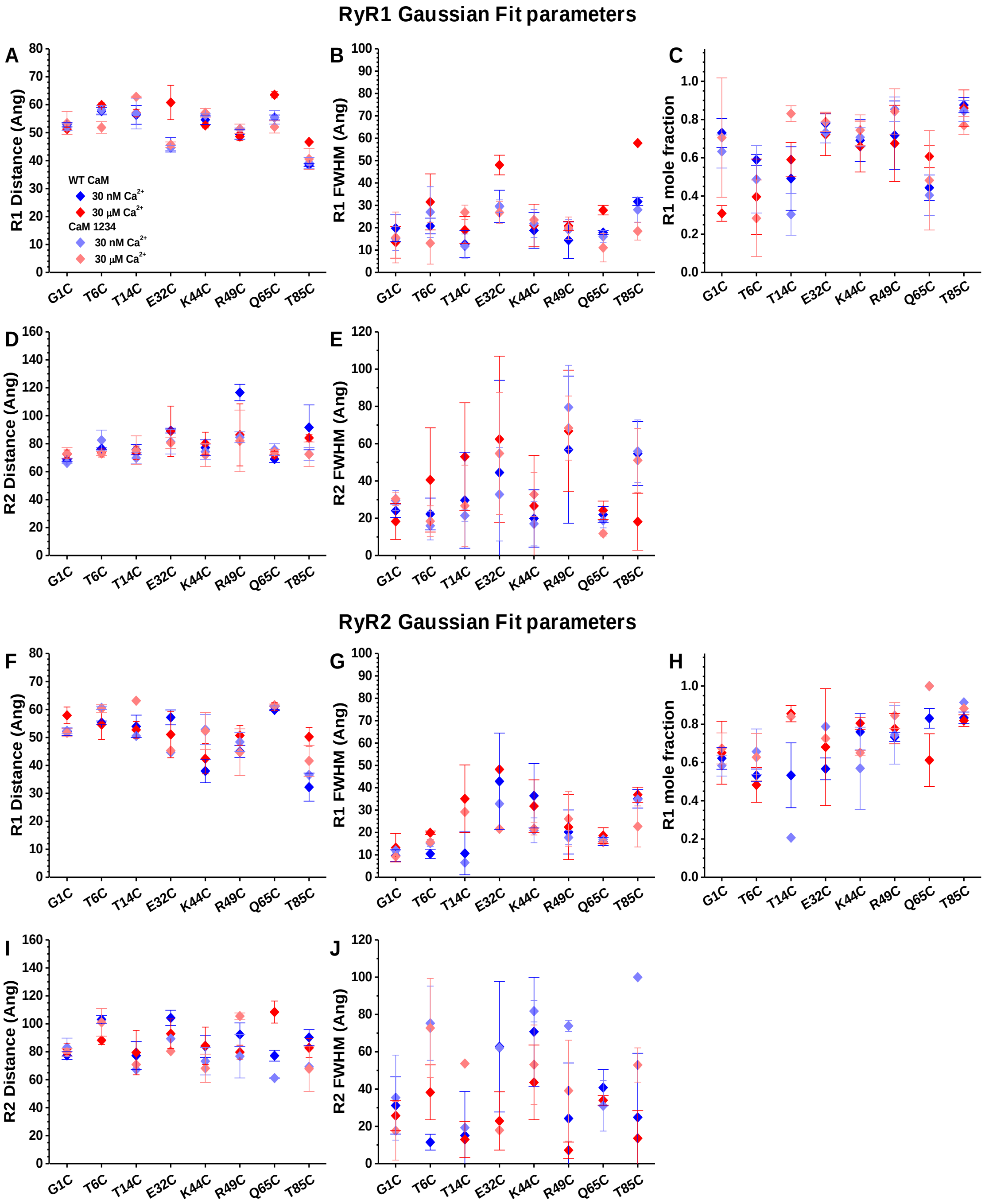


**Supplementary Fig. 3 Gaussian fit parameters for AF568-T26C-CaM.** SR membranes from porcine skeletal (A-E) or cardiac (F-J) muscle were labeled with D-FKBP (AF488-X-FKBP; X = Cys mutation site for fluorescence labeling), and then incubated with 800 nM CaM labeled with acceptor probe at the N-lobe residue T26C. FRET assays used CaM (dark) or $CaM_{1234}$ (light) forms of CaM, and contained 30 nM Ca (blue symbols) or 30 µM Ca (red symbols). Multiexponential analysis of FLT-FRET data yielded a two–distance Gaussian distribution model for the separation between D-FKBP and A-CaM within RyR. Data shown as mean±SEM for the following parameters A and F) shorter distance (R1), B and G) full-width half max (FWHM) for R1, C and H) mole fraction occupied by R1, D and I) longer distance (R2), and E and J) FWHM for R2. N = X-X.

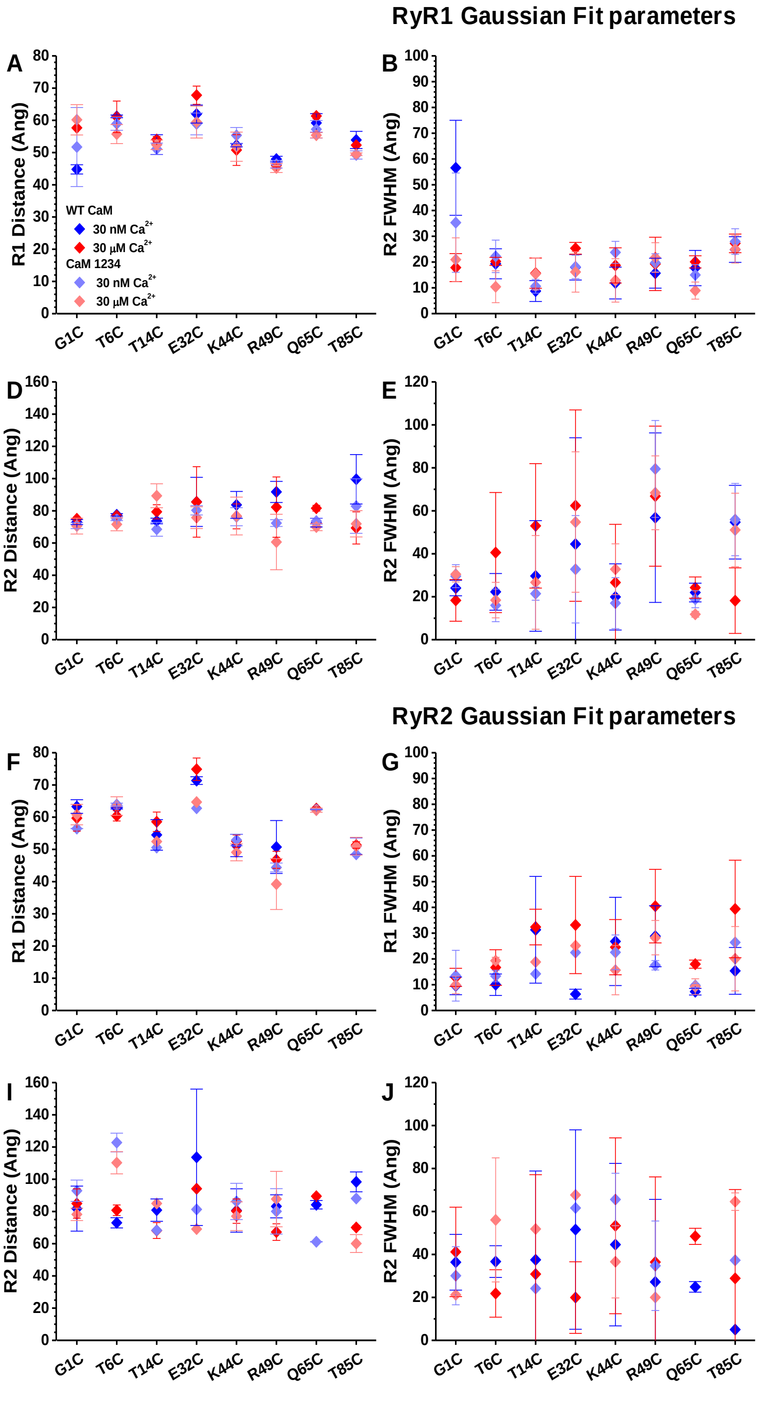


**Supplementary Fig. 4 Gaussian fit parameters for AF568-T34C-CaM.** SR membranes from porcine skeletal (A-E) or cardiac (F-J) muscle were labeled with D-FKBP (AF488-X-FKBP; X = Cys mutation site for fluorescence labeling), and then incubated with 800 nM CaM labeled with acceptor probe at the N-lobe residue T34C. FRET assays used CaM (dark) or $CaM_{1234}$ (light) forms of CaM, and contained 30 nM Ca (blue symbols) or 30 μM Ca (red symbols). Multiexponential analysis of FLT-FRET data yielded a two–distance Gaussian distribution model for the separation between D-FKBP and A-CaM within RyR. Data shown as mean±SEM for the following parameters A and F) shorter distance (R1), B and G) full-width half max (FWHM) for R1, C and H) mole fraction occupied by R1, D and I) longer distance (R2), and E and J) FWHM for R2. N = X-X.

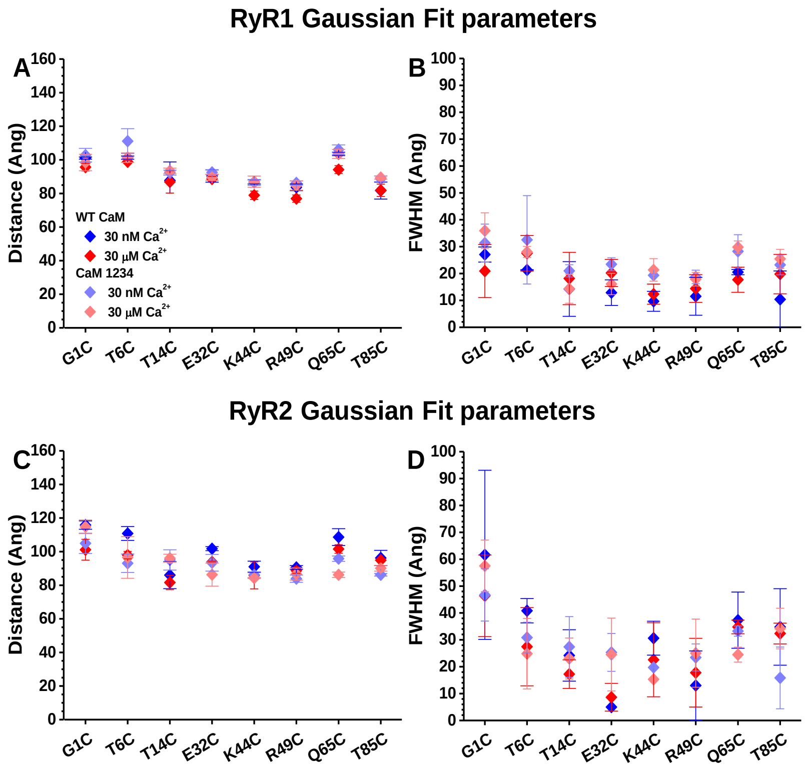


**Supplementary Fig. 5 Gaussian fit parameters for AF568-Y99C-CaM.** SR membranes from porcine skeletal (A and B) or cardiac (C and D) muscle were labeled with D-FKBP (AF488-X-FKBP; X = Cys mutation site for fluorescence labeling), and then incubated with 800 nM CaM labeled with acceptor probe at the N-lobe residue Y99C. FRET assays used CaM (dark) or $CaM_{1234}$ (light) forms of CaM, and contained 30 nM Ca (blue symbols) or 30 µM Ca (red symbols). Multiexponential analysis of FLT-FRET data yielded a one–distance Gaussian distribution model for the separation between D-FKBP and A-CaM within RyR. Data shown as mean±SEM for A and C) distance, and B and D) full-width half max (FWHM) for distance. N = X-X.

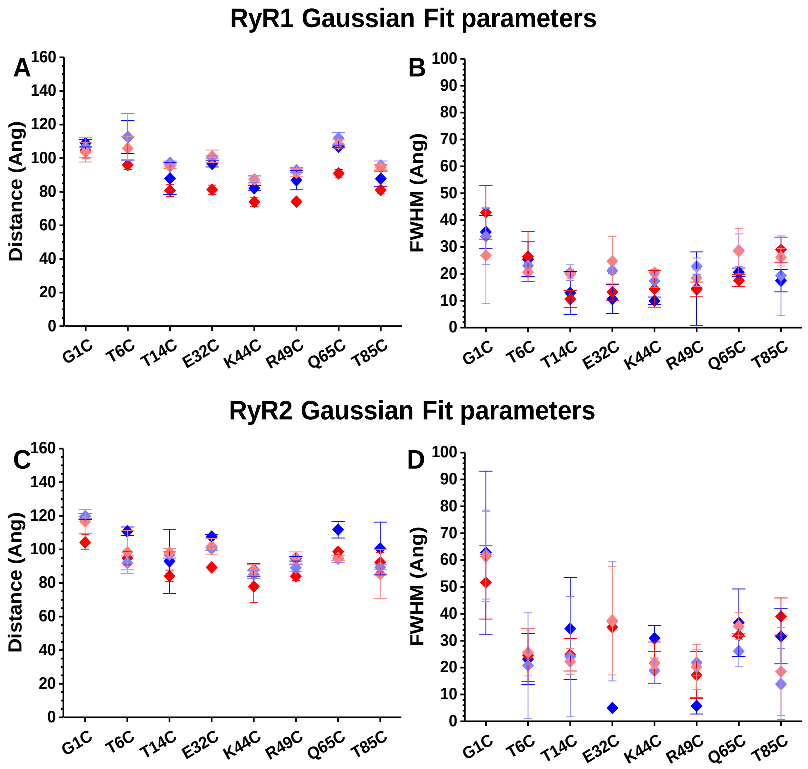


**Supplementary Fig. 6 Gaussian fit parameters for AF568-T110C-CaM.** SR membranes from porcine skeletal (A and B) or cardiac (C and D) muscle were labeled with D-FKBP (AF488-X-FKBP; X = Cys mutation site for fluorescence labeling), and then incubated with 800 nM CaM labeled with acceptor probe at the N-lobe residue T110C. FRET assays used CaM (dark) or $CaM_{1234}$ (light) forms of CaM, and contained 30 nM Ca (blue symbols) or 30 µM Ca (red symbols). Multiexponential analysis of FLT-FRET data yielded a one–distance Gaussian distribution model for the separation between D-FKBP and A-CaM within RyR. Data shown as mean±SEM for A and C) distance, and B and D) full-width half max (FWHM) for distance. N = X-X.

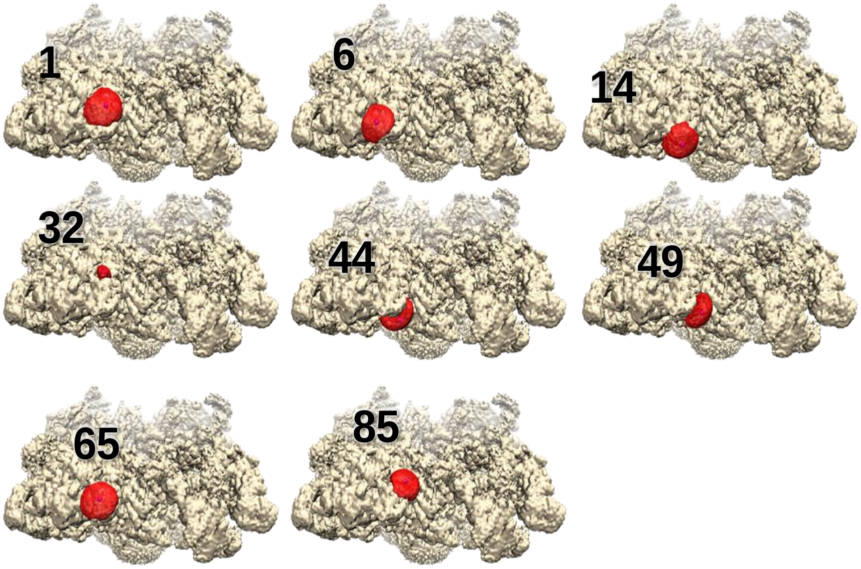


**Supplementary Fig. 7 Donor probe location on FKBP bound to RyR1, determined by simulated annealing.** The cryo-EM density map of RyR1[1] is shown in tan color. The volume sampled by AF488 attached at specific FKBP sequence positions, as indicated, is shown in red. The calculated average position of the probes that is used as the effective probe location in the trilateration calculations are show as purple spheres within the red volumes. In all cases the probe samples a large volume to allow isotropic motion, except for the probe at position 32 where the space is more restricted.

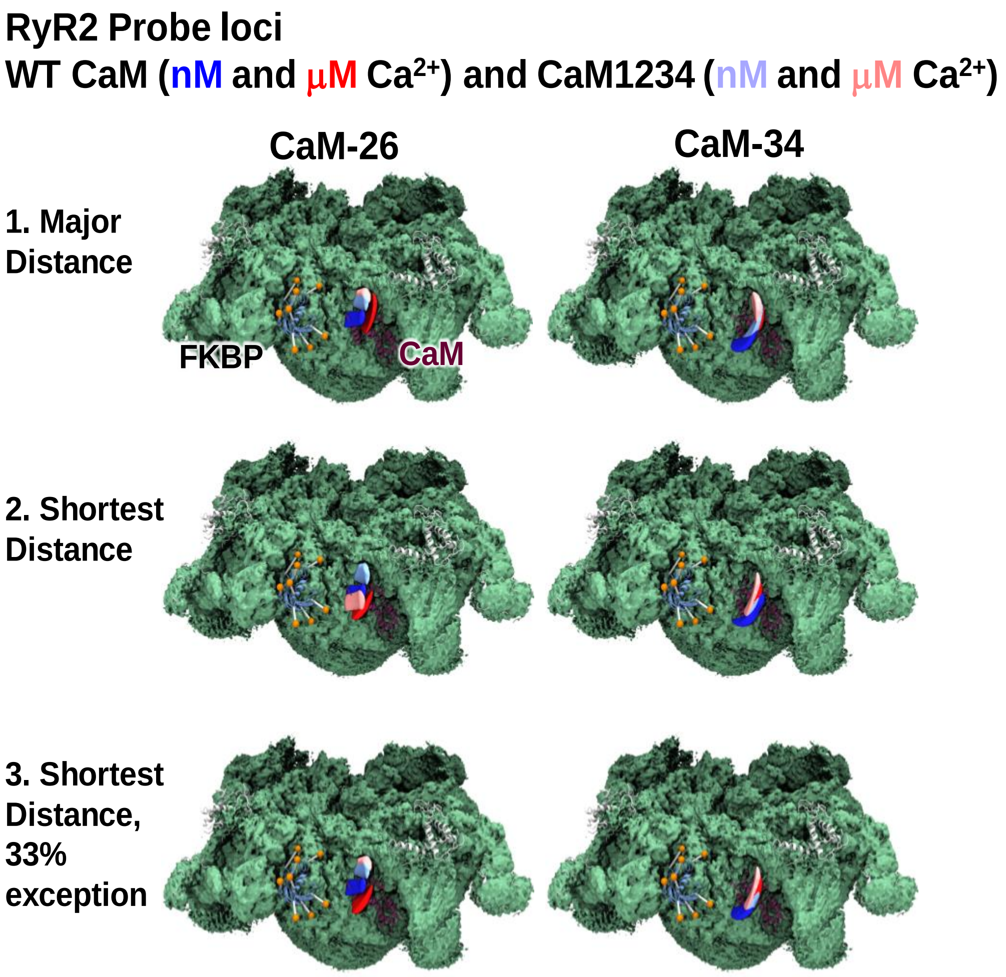


**Supplementary Fig. 8 Acceptor probe loci on RyR2 based on using different criteria for distance selection.** From the two-distance Gaussian fit to the FRET data, acceptor probe location could be selected from one of two distances. Three different sets of trilateration maps were generated based on criteria for selecting distances that were used for trilateration. This included trilateration with 1. the distance with the highest mole fraction (major distance), 2. the shortest distance, and 3. the shortest distance with the exception for when the longer distance had a mole fraction >66%. The latter method was selected as it avoided the case when two distance populations had about equal weight and either the longer or shorter distance was picked in an arbitrary manner. Depending on which selection one uses a difference in the loci are observed. However, method 3 gave the most consistent result with the $CaM_{1234}$ at low and high $Ca^{2+}$ overlapping in location. The cryo-EM density map of RyR2[4] is shown in green. FKBP is shown as ribbon representation in blue with the donor probe positions shown as orange spheres. Trilaterated loci for AF568 probes bound to indicated CaM residues in assay conditions containing 30 nM and 30 µM free $Ca^{2+}$ are blue and red, respectively. Trilaterated loci for probes bound to $Ca^{2+}$ insensitive CaM ($CaM_{1234}$) in assay conditions containing 30 nM and 30 µM free $Ca^{2+}$ are light blue and light red, respectively. The maps of RyR2 are shown at 60° from the cytoplasmic face.

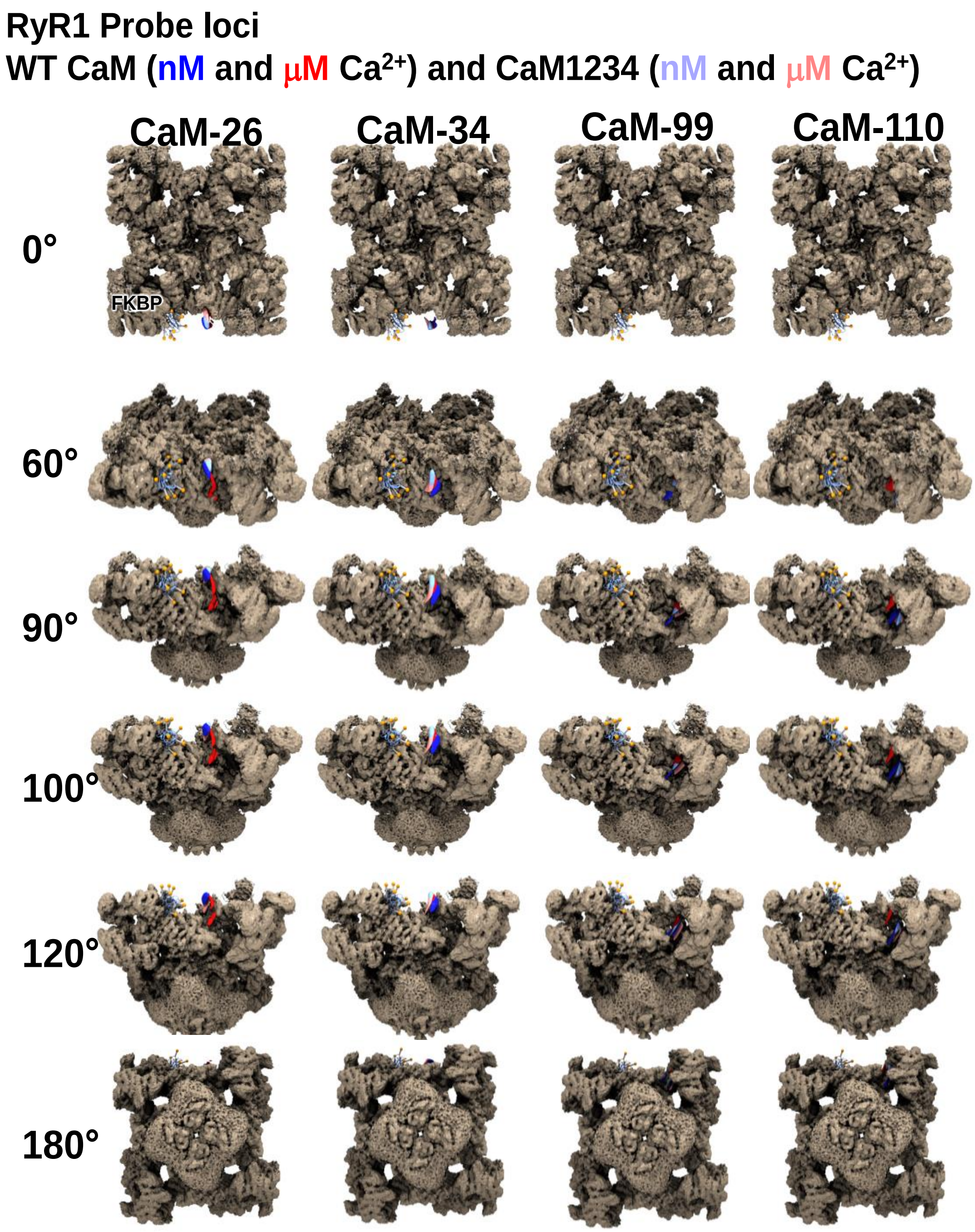


**Supplementary Fig. 9 Acceptor probe loci bound to CaM at 26, 34, 99 and 110 on RyR1.** The cryo-EM density map of RyR1[1] is shown in brown. FKBP is shown as ribbon representation in blue with the donor probe positions shown as orange spheres. Trilaterated loci for AF568 probes bound to indicated CaM residues in assay conditions containing 30 nM and 30 μM free $Ca^{2+}$ are blue and red, respectively. Trilaterated loci for probes bound to $Ca^{2+}$ insensitive CaM ($CaM_{1234}$) in assay conditions containing 30 nM and 30 μM free $Ca^{2+}$ are light blue and light red, respectively. The maps of RyR are shown at indicated degree rotated from the membrane plane.

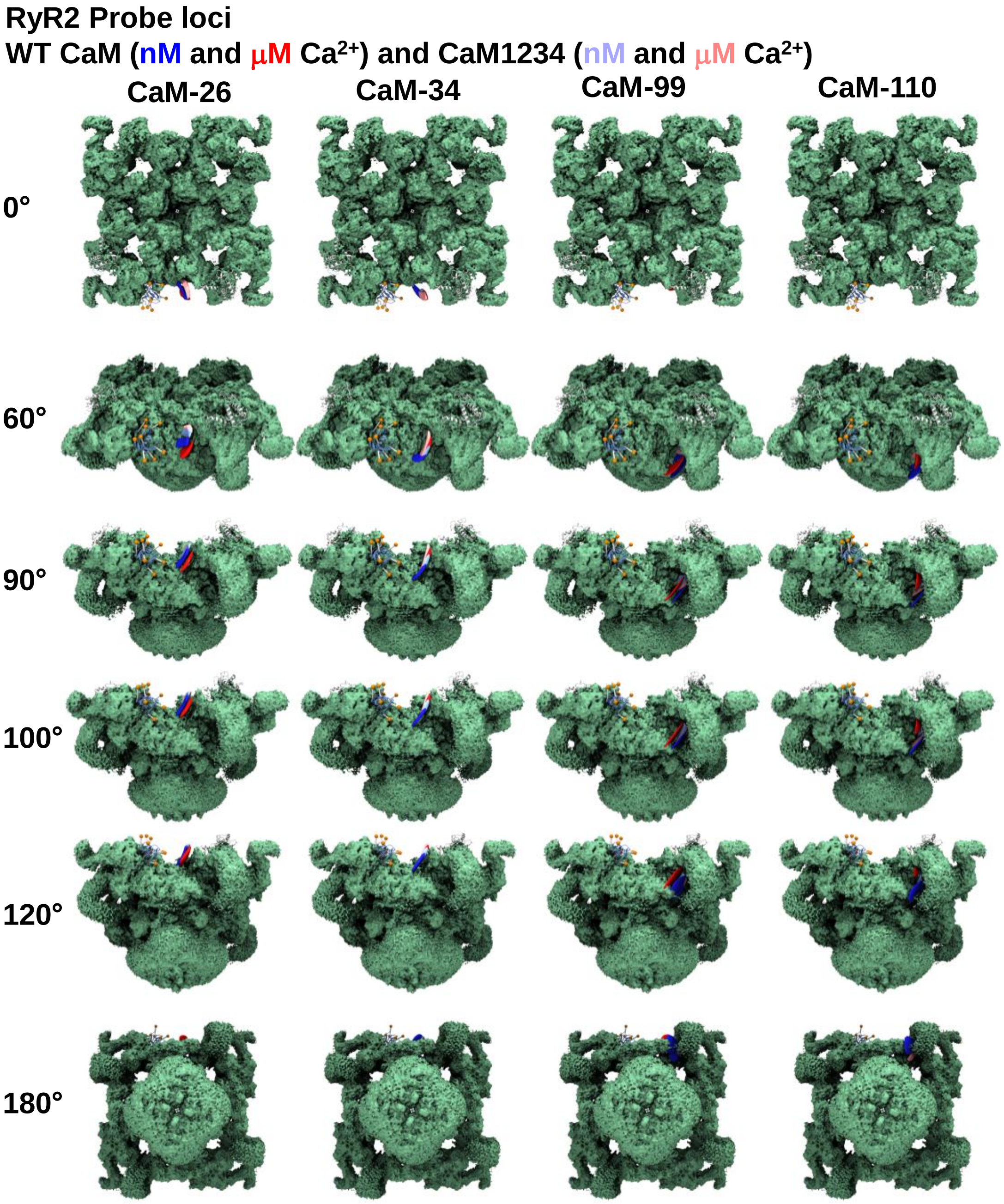


**Supplementary Fig. 10 Acceptor probe loci bound to CaM at 26, 34, 99 and 110 on RyR2.** The cryo-EM density map of RyR2[4] is shown in green. FKBP is shown as ribbon representation in blue with the donor probe positions shown as orange spheres. Trilaterated loci for AF568 probes bound to indicated CaM residues in assay conditions containing 30 nM and 30 μM free $Ca^{2+}$ are blue and red, respectively. Trilaterated loci for probes bound to $Ca^{2+}$ insensitive CaM ($CaM_{1234}$) in assay conditions containing 30 nM and 30 μM free $Ca^{2+}$ are light blue and light red, respectively. The maps of RyR are shown at indicated degree rotated from the membrane plane.

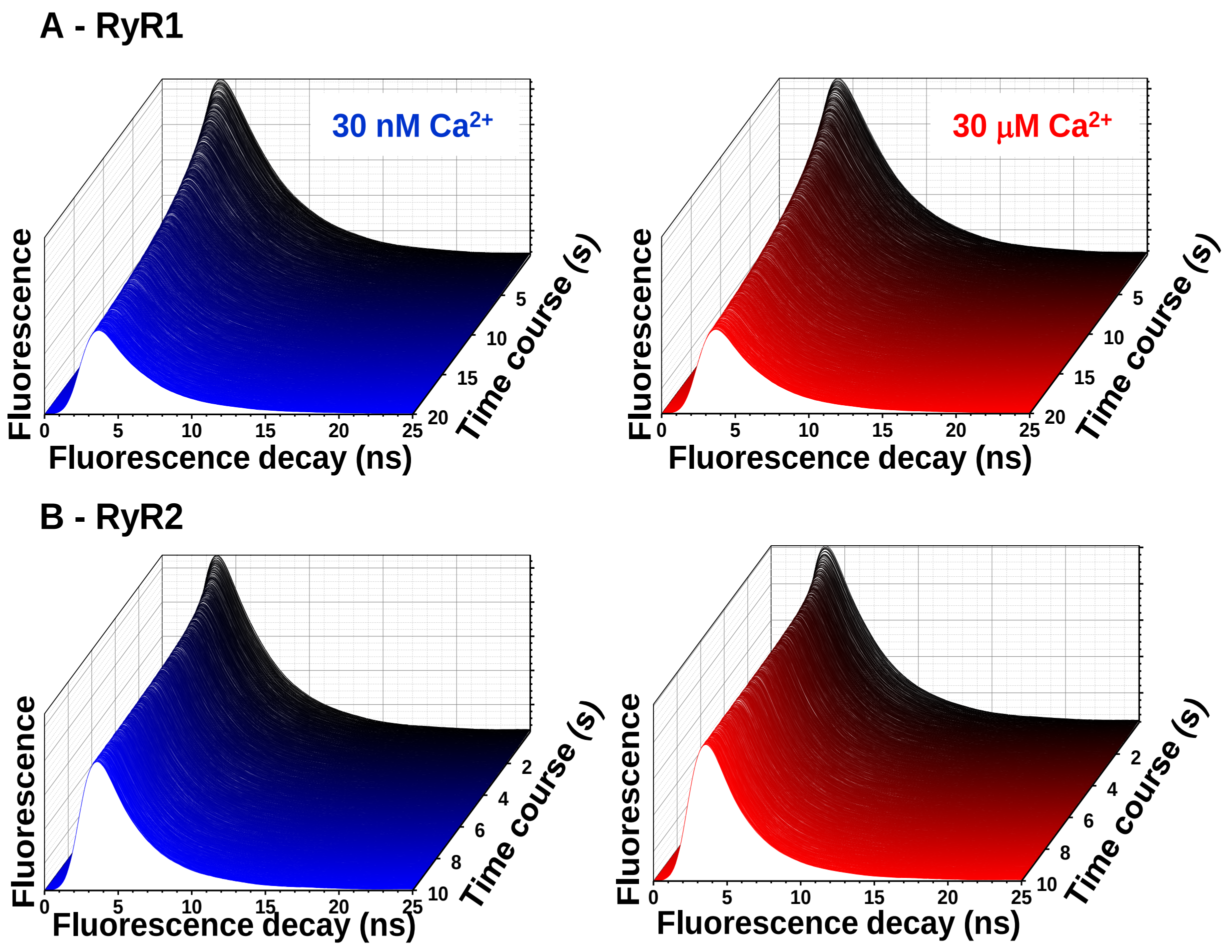


**Supplementary Fig. 11 Representative FLT waveform time course for CaM binding to RyR1 and RyR2 in nM and μM $Ca^{2+}$.** SR membranes from porcine skeletal (A) or cardiac (B) muscle were labeled with D-FKBP (AF488-85-FKBP), and then FLT time course was acquired after rapid (2 ms) mixing with 800 nM (final) A-CaM (AF568-26-CaM). Each graph is a representative FLT time course following mixing. Assay conditions were at 30 nM (blue) or 30 μM (red) $Ca^{2+}$.

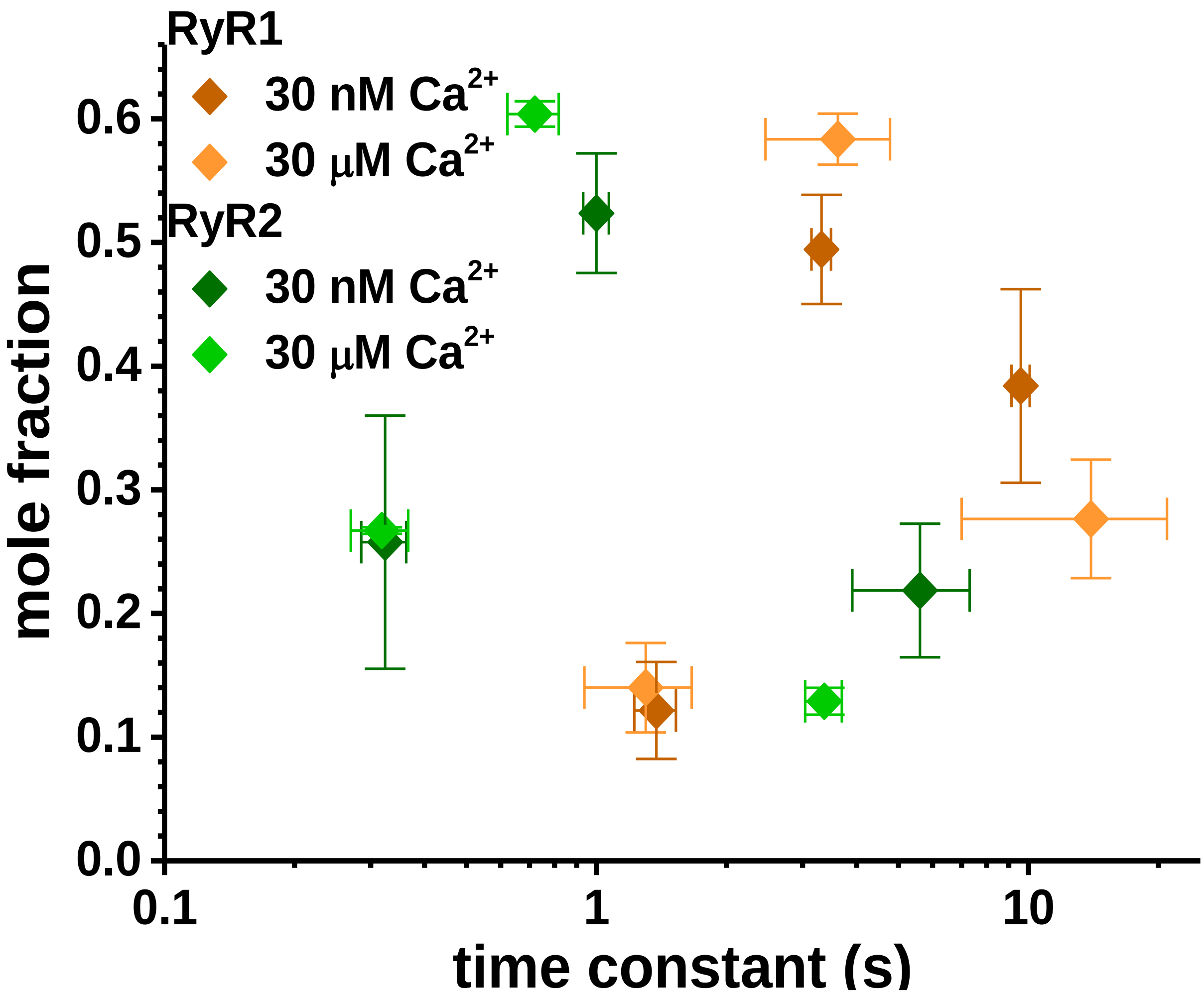


**Supplementary Fig. 12 Three components for nM to μM Ca HSR.** SR membranes from porcine skeletal (brown) or cardiac (green) muscle were labeled with D-FKBP (AF488-85-FKBP), and then FLT time course was acquired after rapid (2ms) mixing with 800 nM (final) A-CaM (AF568-26-CaM. Representative FLT time course are shown in Supplementary Fig X. The FRET data fit best to three-exponential analysis. The parameters of that fit are shown as time constant vs amplitude fraction for CaM binding to RyR1 or RyR2 at 30nM or 30μM $Ca^{2+}$. Data shown as mean±SEM, n = 3.

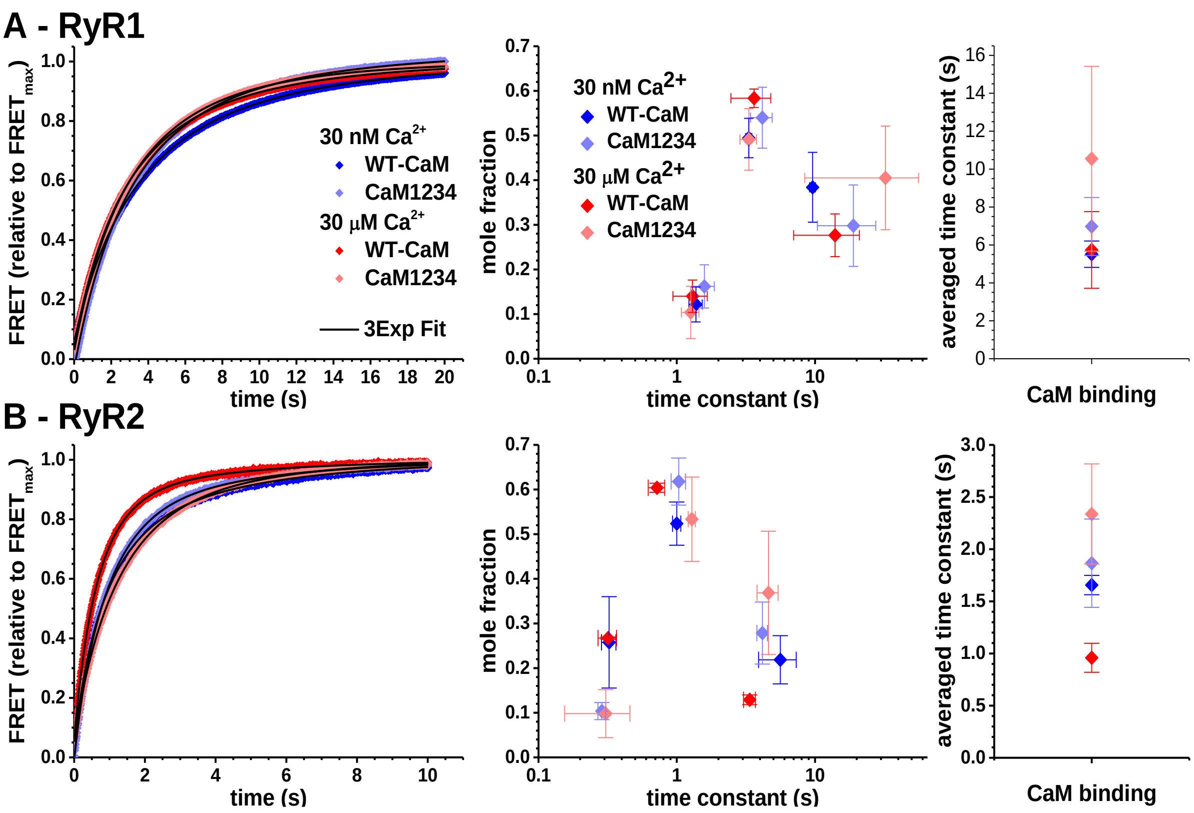


**Supplementary Fig. 13 Kinetics of A-CaM binding to RyR1 and RyR2 in 30 nM and μM $Ca^{2+}$.** SR membranes from porcine skeletal (A) or cardiac (B) muscle were labeled with D-FKBP (AF488-85-FKBP), and then FLT time course was acquired after rapid (2 ms) mixing with 800 nM (final) A-CaM (AF568-26-CaM). Representative FLT time courses shown in Supplementary Fig X. Representative FLT FRET time course following mixing (left panels). All data was fit with three-exponential analysis, with time constant and amplitude values shown in middle panel. In right panel, amplitude-weighted average time constant values for the binding of CaM (dark) or $CaM_{1234}$ (light) to RyR1 (A) or RyR2 (B) at 30 nM (blue) or 30μM (red) $Ca^{2+}$. Data shown as mean±SEM, n = 3.

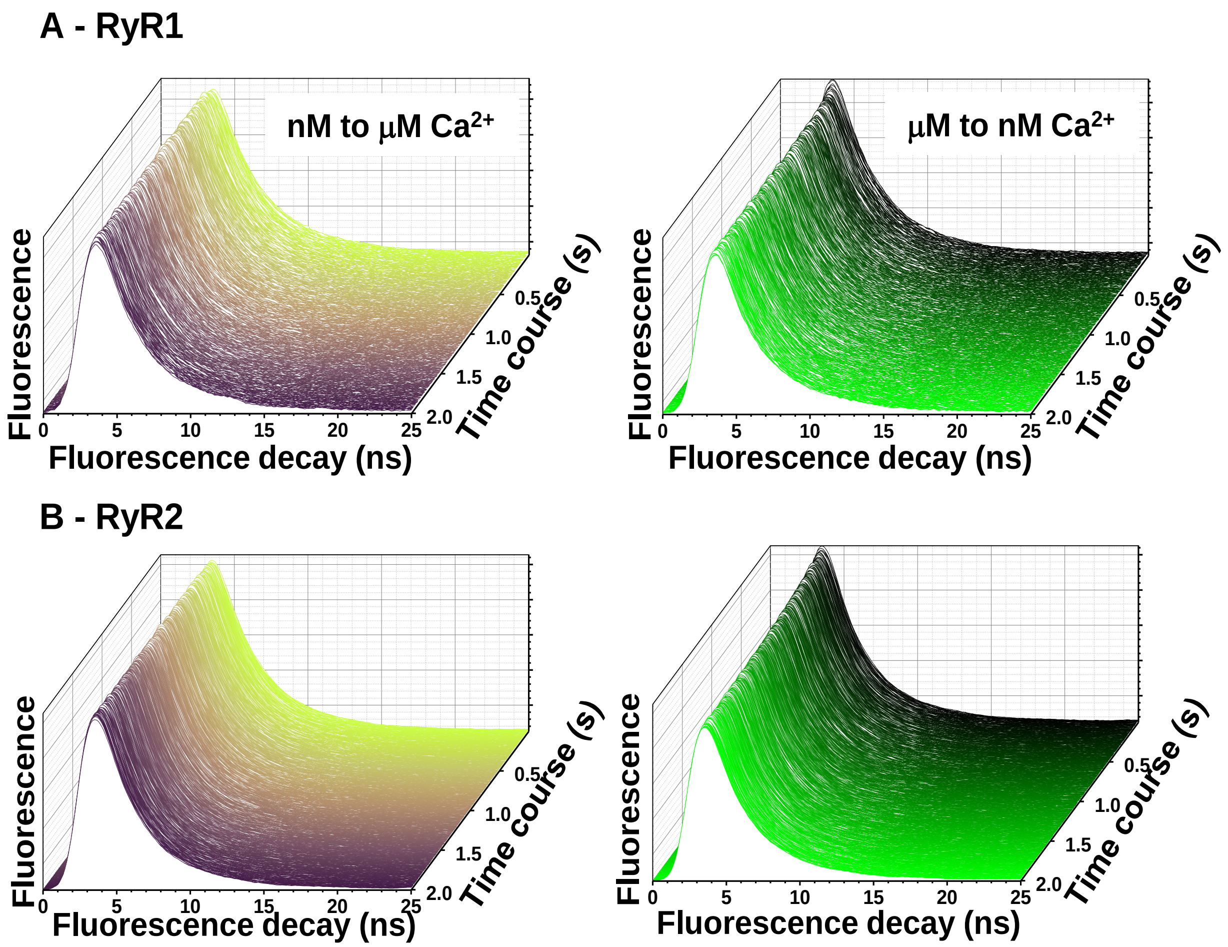


**Supplementary Fig. 14 Representative FLT traces of $Ca^{2+}$ driven RyR-bound CaM structural transitions.** SR membranes from porcine skeletal (A) or cardiac (B) muscle were labeled with D-FKBP (AF488-85-FKBP), incubated with 1.6 μM CaM and then FLT time course was acquired after rapid (2ms) mixing with left panel) $Ca^{2+}$ to increase [$Ca^{2+}$] from 30 nM to μM or right panel) EGTA to reduce [$Ca^{2+}$] from 30 μM to nM. Representative FLT time course following mixing.

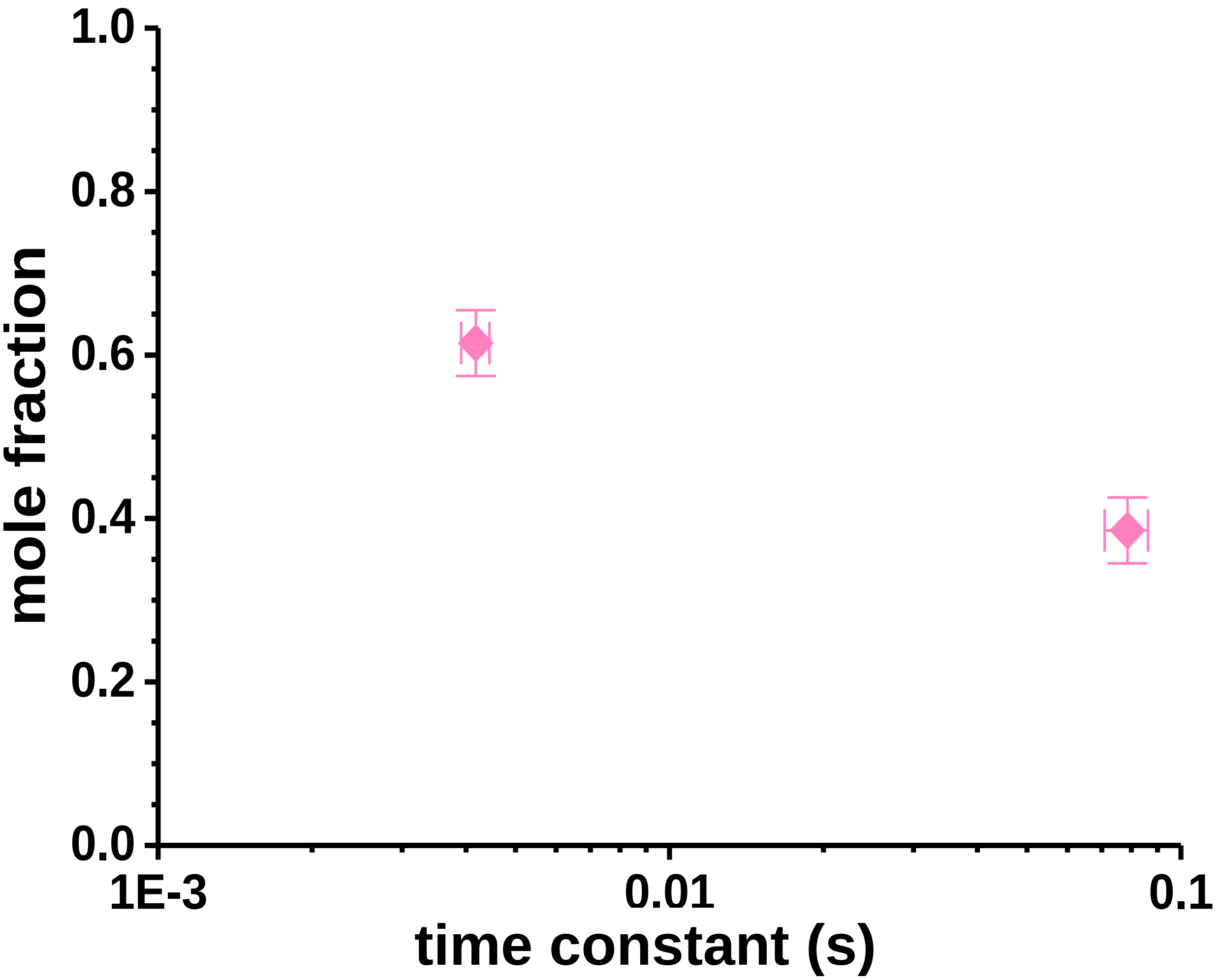


**Supplementary Fig. 15 Parameters of two-exponential fit to FLT FRET data for nM to μM $Ca^{2+}$ driven structural shift of CaM bound to RyR1.** SR membranes from porcine skeletal muscle were labeled with D-FKBP (AF488-85-FKBP), incubated with 1.6 μM CaM and then FLT time course was acquired after rapid (2ms) mixing with $Ca^{2+}$ to increase [$Ca^{2+}$] from 30 nM to μM. Representative FLT waveforms shown in Supplementary Fig 14, and representative FLT-FRET shown in Fig 6. Data fit best to two exponential analysis. Graph displays fit parameters as time constant and amplitude fraction. Data shown as mean±SEM, n = 3.
